# A minimal two-residue C-terminal truncation of AID in adult-onset hyper-IgM syndrome type 2 (HIGM2)

**DOI:** 10.64898/2026.10.27.762264

**Authors:** Kenneth Green, Anne Bruun Rovsing, Ewa Terczyńska-Dyla, Lisbeth Jensen, Karolina Machalica, Jonas Holst Wolff, Lisa Westerberg, Carsten Schade Larsen, Lin Lin, Jacob Giehm Mikkelsen, Jens Magnus Bernth Jensen, Søren Egedal Degn

## Abstract

Activation-induced cytidine deaminase (AID), encoded by *AICDA*, facilitates class-switch recombination (CSR) and somatic hypermutation (SHM) in activated B cells. Loss of AID function causes hyper-IgM syndrome type 2 (HIGM2). While typically autosomal recessive, three C-terminal truncation variants have been reported to act in autosomal dominant manner. We describe a novel heterozygous *AICDA* nonsense variant, c.589G>T (p.G197*), that truncates the final two C-terminal residues of AID and results in adult-onset HIGM2. Single-cell sequencing of primary patient B cells revealed a complete block in CSR and a ∼10-fold reduction in SHM. *AICDA* mRNA expression was balanced between wild-type and variant alleles, yet AID protein abundance was reduced to ∼56% with normal subcellular localization. Remarkably, inducible germinal center (iGB) cultures restored CSR in purified naïve patient B cells. CRISPR-Cas9 editing of healthy donor B cells confirmed that the iGB platform can rescue B cells harboring heterozygous, but not homozygous, *AICDA* variants. These results demonstrate that even a two-amino-acid truncation at the AID C-terminus is sufficient to disrupt its function and cause HIGM2.

## Introduction

Antibodies are central to the adaptive immune response. B cells generate their primary antibody diversity through V(D)J recombination, which assembles immunoglobulin (Ig) gene segments to produce a clonally unique antigen-binding receptor that associates with CD79A/CD79B to form the B-cell receptor (BCR). The same Ig loci later support plasma cell function through expression of secretory heavy-chain isoforms, generating soluble antibodies. Upon binding to cognate antigen, B cells may undergo secondary diversification to acquire new effector functions and improve antigen affinity. This occurs through class-switch recombination (CSR), which replaces the Ig constant region to change antibody isotype (i.e., IgD and IgM to IgG, IgA or IgE), and through somatic hypermutation (SHM), which introduces point mutations into the Ig variable (V) region. Both CSR and SHM are initiated by activation-induced cytidine deaminase (AID) **[1]**.

CSR is achieved by AID-mediated DNA double-strand breaks (DSBs) at donor and acceptor switch (S) regions, followed by non-homologous end joining (NHEJ). CSR is tightly regulated by chromatin architecture and transcription through S regions, including the influence of the 3′ regulatory region (3′RR) and the recruitment of DNA repair factors to S-S junctions **[2–4]**. In SHM, AID deaminates cytosines within the IgV region, generating U:G mismatches. Since uracil pairs with adenine, these mismatches can lead directly to dC-to-dT transitions, if replicated before repair **[5–7]**. While AID directly induces such transitions, the mutational spectrum is expanded through error-prone base excision and mismatch repair pathways. This process is the hallmark of germinal center (GC) affinity maturation, where GC B cell clones undergo iterative cycles of clonal expansion with SHM and subsequent selection for BCR affinity to antigen **[8, 9]**. Beyond affinity improvement, SHM can broaden reactivity through epitope spreading and the emergence of *de novo* specificities from the primary repertoire **[10]**.

The essential role of AID in human CSR and SHM became evident when *AICDA* loss-of-function variants were described in Hyper-IgM Syndrome Type 2 (HIGM2) patients **[11]**. HIGM2 is a B-cell-intrinsic inborn error of immunity characterized by normal to elevated serum IgM, absent switched isotypes (IgG/IgA/IgE), and recurrent respiratory infections **[12, 13]**. Autosomal-recessive (AR) AID deficiency usually presents in early childhood with recurrent sinopulmonary infections, absent switched memory B cells, and complete loss of both CSR and SHM **[12, 14]**. In contrast, the three reported heterozygous C-terminal truncating variants (p.V186*, p.L189*, p.R190*) cause autosomal-dominant (AD) disease **[15–17]** with widely varying severity. Symptoms range from severe antibody deficiency with bronchiectasis to carriers without recurrent infection identified only by family screening **[18–20]**.

These AD HIGM2 variants have been particularly informative because they differentiate CSR from SHM. Functionally, the N-terminal catalytic domain of AID is sufficient to generate SHM, whereas CSR additionally requires the C-terminus **[21–25]**. Mechanistically, the C-terminus couples AID-mediated DNA cleavage with the S-S joining process by recruiting DNA repair and end-protection factors such as UNG, MSH2/6, 53BP1 and DNA-PKcs, as well as heterogeneous nuclear ribonucleoproteins (hnRNPs) **[3, 22, 25–28]**. The C-terminus has also been shown to regulate AID oligomerization, which influences which hnRNP cofactors are recruited: monomeric AID favors hnRNP-K for DNA cleavage, while dimeric AID recruits hnRNP-L/U/SERBP1 to promote recombination **[4, 28, 29]**. Beyond its role in S-S joining, the C-terminus contains the canonical nuclear export signal (NES) and a eukaryotic translation elongation factor 1-alpha 1 (eEF1A1)-dependent cytoplasmic retention motif, together keeping >90% of AID in the cytoplasm **[30–34]**.

The dominant-negative behavior of C-terminal truncating variants remains mechanistically unresolved. Truncated AID has been reported to heterodimerize with the wild-type protein **[22]**, to sequester shared cofactors **[35]**, to form nuclear condensates that withdraw wild-type AID from the IgH locus **[36]**, or to interfere with the repair of AID-induced lesions in a manner dependent on catalytic activity rather than abundance **[26, 37]**. Alternatively, the dominant phenotype in these cases may stem simply from haploinsufficiency, a reduction in functional AID dosage alone. Yet, this fails to explain the near-normal CSR and SHM reported in asymptomatic carriers of *AICDA* null alleles **[38, 39]**.

Here, we describe a previously unreported heterozygous AID variant, p.G197*, that truncates just the last two C-terminal residues yet causes adult-onset HIGM2. The patient’s B cells exhibited complete CSR failure and a ∼10-fold reduction in SHM. We used inducible GC B cell (iGB) cultures to activate the patient’s naïve B cells and observed that while *AICDA* was expressed from both alleles and subcellular localization remained unaffected, total AID protein level was reduced to ∼56% compared to healthy donor cells. Furthermore, using the iGB cultures, we observed unexpectedly that the patient’s naïve B cells underwent CSR, albeit less efficiently than controls. Likewise, CRISPR-Cas9 deletion of the AID C-terminus in healthy donor B cells, creating engineered heterozygous C-terminal truncated cells, also retained CSR in iGB cultures. These findings show that sustained activation can restore CSR despite lower AID levels, suggesting that the dominant-negative effects of C-terminal truncations depend on the B cell’s activation state.

## Results

### Hyper-IgM syndrome type 2 (HIGM2) patient phenotype and genotype

A 34-year-old Danish female, a former elite athlete, presented with 4-5 years of recurrent upper respiratory tract infections, intermittent bacterial sinusitis, and a recent pneumonia. Infections in childhood were unremarkable apart from an appendectomy at 6 years and thoracic shingles at 15 years of age. She was initially referred for suspected asthma, which was ruled out by a negative methacholine challenge. Early high-resolution chest CT showed mild bilateral bronchiectasis. There was no family history of primary immunodeficiency or recurrent/unusual infections.

Serum immunoglobulins demonstrated a hyper-IgM phenotype with elevated IgM (4.61 g/L) and undetectable IgG (<0.8 g/L) and IgA (<0.08 g/L) (**Fig. 1A**). No M-component was detected in serum or urine, indicating that the increased IgM was of polyclonal origin. Bone-marrow examination revealed normo-to mildly hypercellular marrow with preserved trilineage hematopoiesis, a normal distribution of polyclonal plasma cells, and no dysplasia, infiltration, fibrosis, amyloid, or clonal plasma-cell expansion. CT of the axial skeleton and hematologic assessment found no evidence of myeloma, MGUS, amyloidosis, or other hematologic malignancy. Peripheral blood counts were normal without cytopenias except for transient thrombocytopenia during earlier pregnancy. Liver enzymes were within the reference range apart from a pregnancy-limited rise in alkaline phosphatase. No lymphadenopathy, splenomegaly, opportunistic infections, autoimmune phenomena, malignancy, or sclerosing cholangitis were identified. Flow cytometry confirmed defective CSR as patient memory B cells (CD27⁺IgD⁻) were reduced and lacked switched BCRs compared with healthy controls (**Fig. 1B–C; Fig. S1A**). Following initiation of IgG-replacement therapy (**Fig. 1D**), the patient has remained free of significant infections.

**Figure 1.**
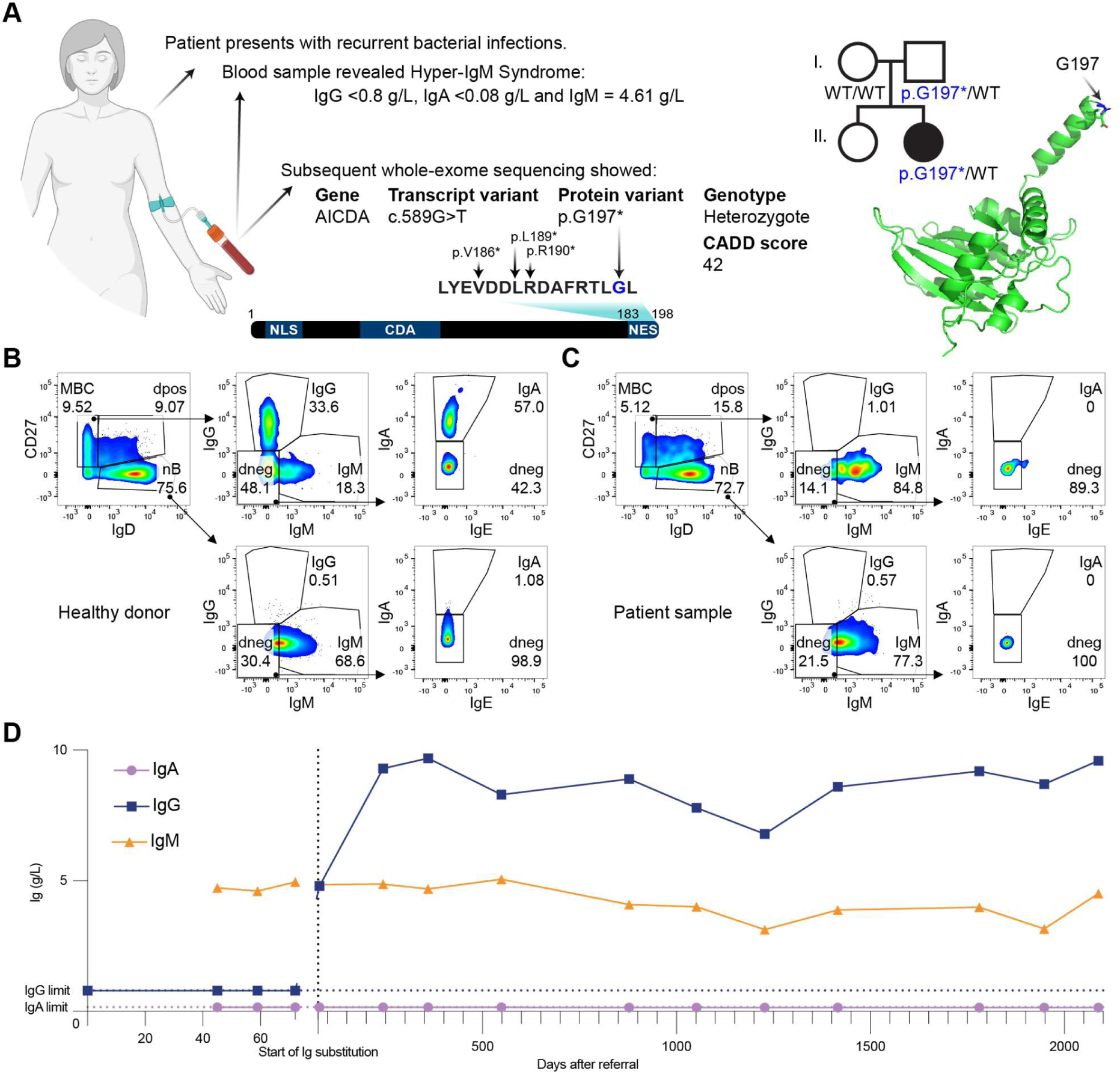
Clinical presentation, genetic diagnosis, and baseline B-cell phenotype. **(A)** Case overview and serology. Summary of clinical details and serum Ig levels showing elevated IgM with undetectable IgG/IgA. Center, genetic findings: TWIST-WES and PCR-free WGS identified heterozygous *AICDA* c.589G>T (p.G197*). Schematic of AID domains highlights the C-terminal region with nuclear export/retention motifs; known autosomal-dominant HIGM2 variants (p.V186*, p.L189*, p.R190*) are mapped, and p.G197 (site of truncation) is highlighted in blue. Right, AlphaFold2 AID model with p.G197 highlighted in blue (AF-Q9GZX7-F1-v6). NES: nuclear export sequence; NLS: nuclear localization signal (NLS); CDA: cytidine deaminase domain. **(B)** Control blood. Representative flow cytometry from peripheral blood: gating on live/singlets/CD20⁺ B cells with intracellular Ig isotype readouts (panel and antibodies in supplementary table 1; full gating tree in Fig. S1A). **(C)** Like (B) but patient blood. Matched gating shows reduced memory (CD27⁺IgD⁻) and absence of switched Igs in patient memory B cells. **(D)** Longitudinal serum-IgM, -IgG and -IgA measurements for a five-year period from patient referral to today. Start of intravenous IgG treatment is marked by a vertical dotted line, and results in IgG-levels within reference (between 6.9-15.7 g/L). IgM was elevated throughout with a measured amount well above the upper reference of 2.3 g/L. Horizontal dotted lined mark detection limit of IgG and IgA, respectively. IgE as well as IgG1-4 have additionally been monitored as being below their detection-limits.

Whole-exome sequencing identified a novel heterozygous nonsense variant in *AICDA*, c.589G>T (p.G197*). The substitution introduces a premature stop codon, truncating the final two C-terminal residues of AID. The variant had a high predicted pathogenicity (CADD 42) and was absent from population and clinical variant databases (gnomAD, Ensembl, ClinVar, TOPMed/BRAVO, dbSNP). Sanger sequencing confirmed the variant in the patient and revealed paternal inheritance (**Fig. 1A**). The father, although a heterozygous carrier of the variant, was clinically well with normal immunoglobulin levels and no history of recurrent infections. The mother carried only the reference allele. Plasma IgA and IgM were within reference intervals in both parents, and IgG was normal in the father and slightly elevated in the mother. No additional variants relevant to humoral immunity were identified by subsequent trio whole-genome sequencing (WGS).

In summary, these findings establish a diagnosis of adult-onset HIGM2 associated with a novel heterozygous *AICDA* truncation (p.G197*). However, a single affected patient and a single unaffected carrier do not establish the mode of inheritance.

### Structural, evolutionary and population-level context of the AID C-terminus

To place p.G197* among known disease-associated variants, we assembled all alleles reported in patients with HIGM2 using PubMind, a large language model framework for extracting variant-disease associations from PubMed, followed by manual curation **(Fig 2A) [40]**. Fifty-five distinct variants have been described, including 31 missense, 13 nonsense, 6 frameshift, 4 in-frame indels, and 1 splice-site change. Fifty-two variants distributed throughout the first 182 residues segregate as AR variants. By contrast, all three AD variants (p.V186*, p.L189*, p.R190*) are confined to the C-terminal region **[15–17]**.

**Figure 2.**
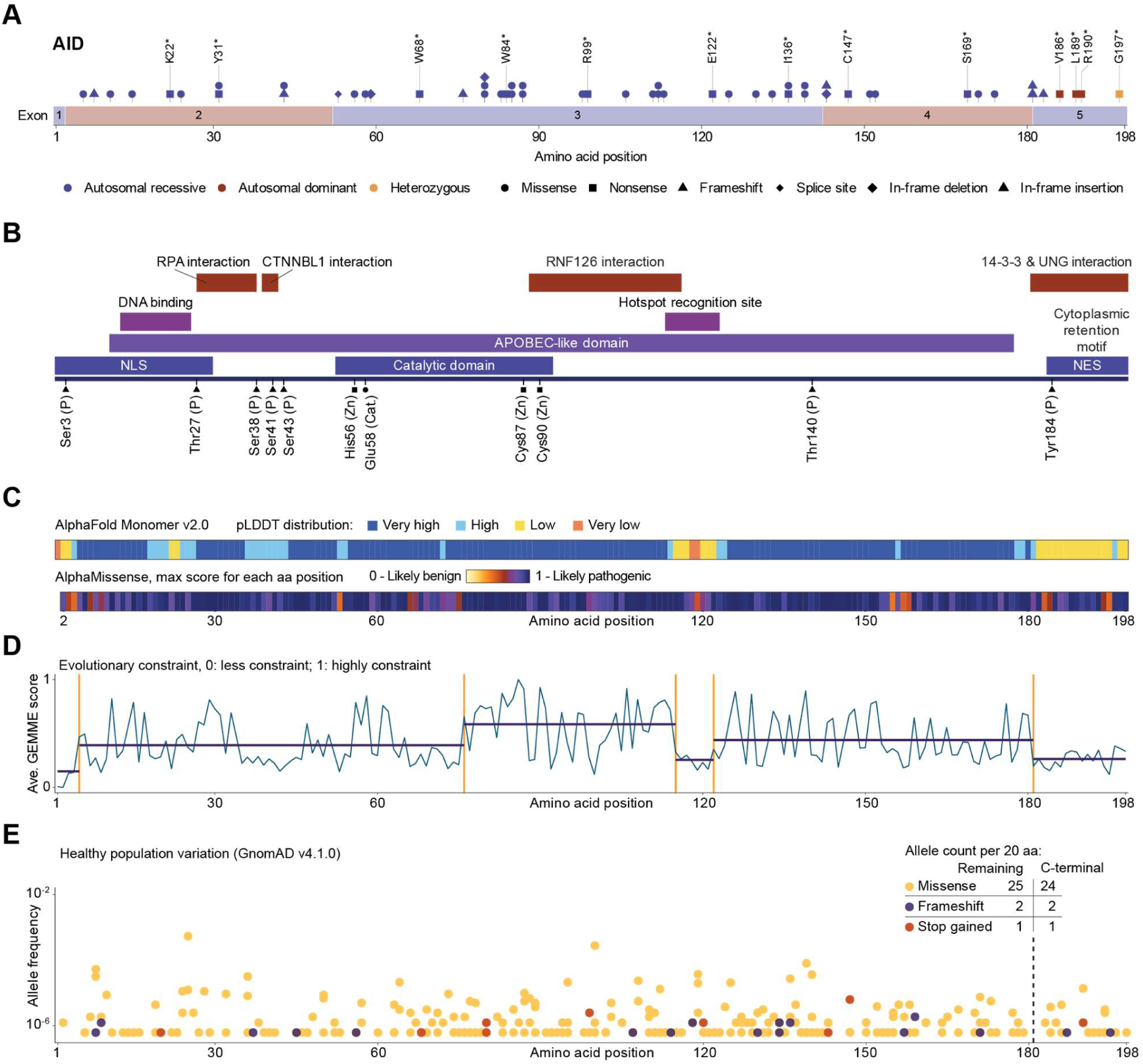
Structural, evolutionary, and population-level analyses of AID. (**A**) HIGM2 patient variant overview retrived using PubMind **[40].** (**B**) Domain architecture of AID annotated from UniProt, phosphosite databases, and published studies **[32, 43, 48]**. The protein encompasses an APOBEC-like deaminase fold with a central catalytic domain, a nuclear localization signal (NLS), a nuclear export signal (NES), and a C-terminal cytoplasmic retention motif. Key functional regions include the DNA-binding domain, hotspot recognition site, and interaction interfaces for RPA, CTNNBL1, RNF126, 14-3-3, and UNG. Phosphorylation sites (P), zinc-coordinating residues (Zn), and the catalytic glutamate (Cat) are indicated below. The patient variant p.G197* maps to the C-terminal 14-3-3/UNG interaction domain. (**C**) AlphaFold2 (Monomer v2.0) predicted structure of AID colored by predicted local distance difference test (pLDDT) scores. Below, per-residue AlphaMissense pathogenicity scores (maximum score per position) are displayed along the protein, with scores ranging from 0 (likely benign) to 1 (likely pathogenic). (**D**) Evolutionary constraint profile displayed as mean per-residue GEMME scores (0, less constrained; 1, highly constrained). Horizontal bars indicate segment-level constraint boundaries inferred by ProteoCast; vertical orange lines denote segment boundaries. (**E**) Distribution of *AICDA* variants from gnomAD v4.1.0 plotted by amino acid position and allele frequency (log_10_ scale). Variants are colored by consequence. Dashed line at position 181 marks the C-terminal segment. Inset table summarizes variant counts normalized per 20 amino acids for the remainder of the protein (residues 1–181) versus the C-terminal region (residues 182–198).

Structurally, AID is organized into specific functional domains that determine its activity, localization, and regulatory interactions. AID contains an APOBEC-like domain that provides the conserved structural scaffold shared across the APOBEC family of cytidine deaminases (**Fig. 2B) [41, 42]**. Within this region, the catalytic domain includes the essential zinc-binding residues (His56, Cys87, Cys90) and catalytic Glu58 that drive C-to-U deamination **[43]**. The APOBEC-like domain includes sites interacting with replication protein A (RPA) and catenin beta like 1 (CTNNBL1) as well as a DNA binding site and hotspot-recognition site, which all mediate ssDNA binding and enforce WRC/GYW motif targeting within Ig loci, thereby influencing deamination efficiency and specificity **[43–47]**. The E3 ligase RNF126 also binds to the APOBEC-like domain and ubiquitylates AID to restrict its active lifetime and limit off-target activity **[48]**. The C-terminal tail interacts with both 14-3-3, which recruits AID to S regions in the Ig locus, and with UNG, which coordinates processing of CSR-specific lesions by removing uracils leading to recruitment of DNA base excision repair proteins **[25, 26, 49]**. Furthermore, the N-terminal contains a nuclear localization signal (NLS) motif, and the C-terminal both a NES and an eEF1A1-dependent cytoplasmic retention motif **[30, 31]**.

AlphaFold2 modelling predicted a well-structured catalytic fold but low predicted Local Distance Difference Test (pLDDT) values across both termini, including the DNA-binding region (aa 13–26) and the hotspot-recognition loop (aa 113–123), indicating that these segments function as intrinsically disordered regions (IDRs) (**Fig. 2C**). This pattern aligned with biochemical studies showing that these flexible regions mediate interactions with DNA, cofactors, and trafficking regulators rather than forming stable secondary structure **[4, 29, 33]**. AlphaMissense scores further highlighted the functional relevance of these disordered segments, as many residues, including within the C-terminal tail received high predicted pathogenicity scores, supporting the idea that short linear motifs embedded within the C-terminal are essential **[50]**.

Using GEMME, a global epistatic model that estimates evolutionary tolerance to amino-acid substitutions, mutational sensitivity profiling across the full AID protein showed strong constraint within the catalytic core and decreased constraint across the N-and C-terminal regions (**Fig. 2D**). Furthermore, alignment of the C-terminal residues across vertebrate species revealed sequence flexibility, including substitutions and length variation (**Fig. S2A**).

Using population-level variation from gnomAD v4.1.0 (807,162 individuals), constraint estimates indicated that *AICDA* is only modestly constrained (synonymous o/e = 1.10; missense o/e = 0.83; pLoF o/e = 0.77, pLI = 0). To assess regional patterns, we compared the distribution of missense, frameshift, and stop-gained variants across the structured portion of AID (aa 1–181) and the C-terminal tail (aa 182–198) and observed nearly identical variant densities between regions (missense: 25 vs 24; frameshift: 2 vs 2; stop-gained: 1 vs 1; **Fig. 2E**). CADD burden was similarly uniform, with comparable numbers of high-scoring variants (CADD > 20: 23 vs 22 per 20 aa; **Fig. S2B**), and ClinVar pathogenic and likely pathogenic variants were distributed across the protein without C-terminal depletion (**Fig. S2C**). Given AID’s small size and the limited mutational target in the C-terminal segment, larger datasets will be required to reliably detect subtle constraint.

Altogether, this suggests that the p.G197* truncation disrupts the NES, the cytoplasmic retention motif, and the UNG and 14-3-3 interaction sites that help direct CSR. Furthermore, these analyses indicate that while the C-terminus lacks clear population-level variation depletion, critical individual residues within this region still carry high predicted pathogenicity scores.

### Somatic hypermutation is present but markedly reduced in patient B cells

To assess SHM, we performed single-cell 5′ RNA-seq with paired BCR sequencing on CD19⁺ B cells from the patient and a healthy control. After preprocessing with Cell Ranger and quality control using Seurat v5 **[51]**, transcriptionally defined B-cell subsets were identified, including naïve, memory, atypical memory, and plasma cells (**Fig. 3A–B**; **Fig. S3A–B**).

**Figure 3.**
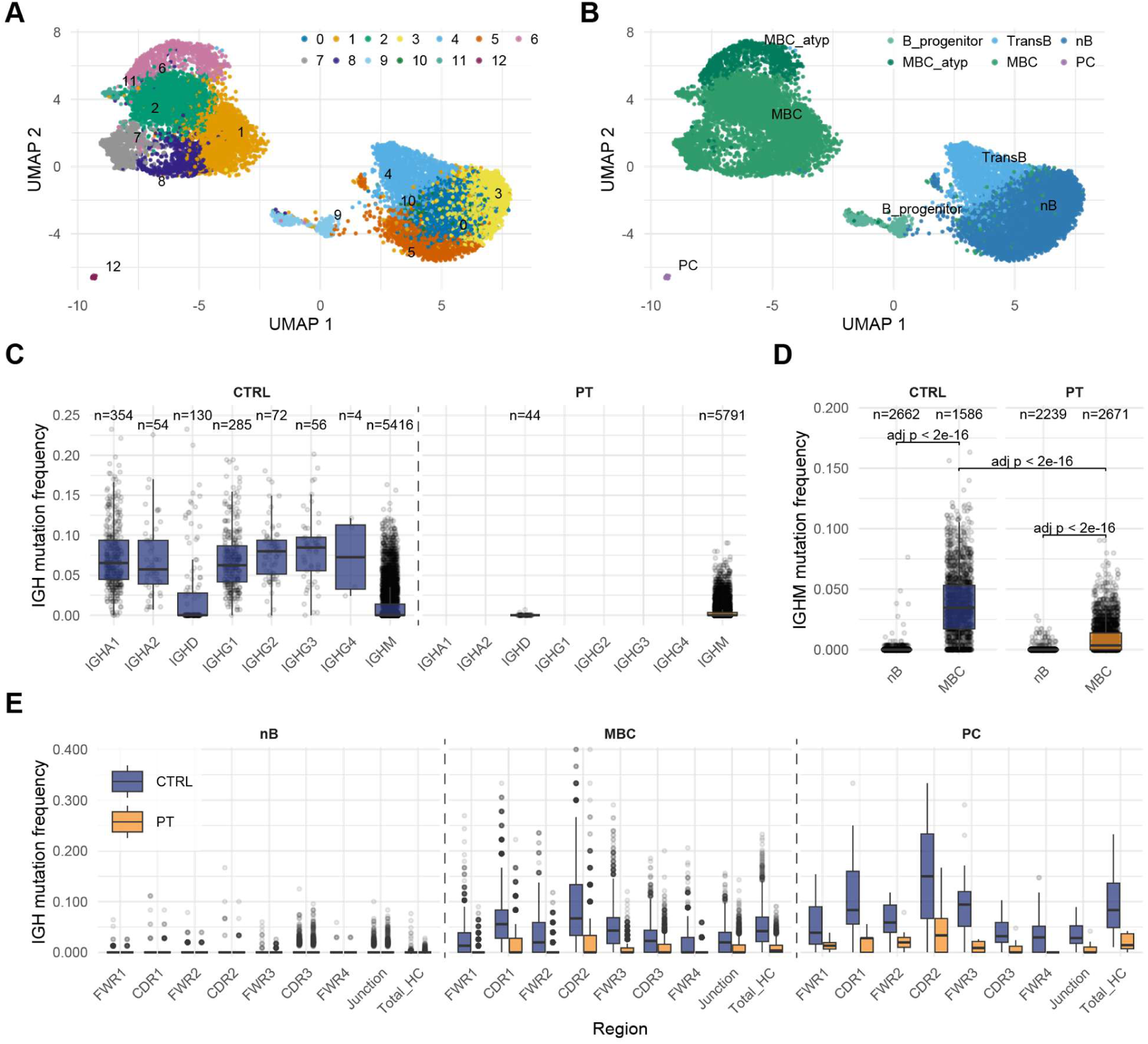
Single-cell transcriptomes and BCR-seq reveal preserved but reduced SHM in patient B cells. **(A)** Joint gene expression UMAP of control (CTRL) and patient (PT) CD19+ B cells after QC, doublet removal, and CCA-based integration, showing 13 clusters. **(B)** (A) with cluster annotations according to major B-cell subtypes (progenitor, transitional, naïve [nB], atypical memory, memory [MBC], plasma cell [PC]). **(C)** Heavy-chain isotype calls from paired V(D)J sequencing; PT lacked class-switched IGH transcripts, corroborating serum/BCR protein data. 6371 and 5835 cells shown for CTRL and PT, respectively (with respectively 19 and 2 cells showing NA for confident isotype call). **(D)** Per-cell IGHM V-region mutation frequency (vs reconstructed germline) in nB and MBC. Both CTRL and PT show significant nB→MBC increases (Hodges–Lehmann [HL] shift; Wilcoxon rank-sum with Benjamini–Hochberg correction, p_adj < 2×10⁻¹⁶). Median SHM in IGHM MBCs was ∼10-fold lower in PT than CTRL (0.00347 vs 0.0347; HL_PT–CTRL = −0.0273; p_adj < 2×10⁻¹⁶). **(E)** Region-resolved IGH SHM in nB, MBC and PC (IMGT segments, FWR1–3, CDR1–3, AIRR junction and IGH total mutation frequency ‘Total_HC’): PT shows clear SHM across regions in MBC and PC, but in a smaller fraction of cells than CTRL. All nB→MBC shifts were significant (p_adj < 2×10⁻¹⁶). BCR-seq analyses included cells with exactly one productive IGH paired to one IGK/L (CTRL n=6,390; PT n=5,837). See Supplementary Statistics 1–2 and 7–9. Group comparisons used two-sided Wilcoxon rank-sum tests with Benjamini–Hochberg FDR; when effect sizes are shown, Hodges– Lehmann median shifts with 95% CIs are reported. CTRL: control; PT: patient.

BCR sequencing was processed with Cell Ranger VDJ and the Immcantation suite **[52]** to assign clones, reconstruct germline V(D)J sequences, and quantify SHM. Only cells with paired *IGH* and a single light chain (*IGK* or *IGL*), and with matching gene-expression profiles, were analyzed (6390 control; 5837 patient). As expected, patient B cells lacked class-switched transcripts, whereas control B cells showed a diverse repertoire of isotypes (**Fig. 3C**; **Fig. S3C**).

To quantify SHM, we measured *IGHM* V-region mutation frequency. Naïve B cells from both patient and control had minimal mutations (median = 0), whereas memory B cells (MBCs) had accumulated mutations. However, patient MBCs carried ∼10-fold fewer *IGHM* mutations than control (median 0.0035 vs. 0.0347; Hodges–Lehmann shift −0.0273; p<2×10^-16^; **Fig. 3D**; **SuppStat 1–2**). This reduction persisted when analyzing all IGH isotypes together (**Fig. S3D**; **SuppStat 3–4**), and when examining Ig light-chain SHM (*IGK/IGL*) (**Fig. S3E**; **SuppStat 5–6**). Thus, SHM occurs in the patient but at markedly reduced levels.

To evaluate mutation targeting, we mapped mutations across complementarity-determining region (CDR)2, framework region (FWR)3, and CDR3. Control MBCs showed expected enrichment of mutations in these regions, whereas the patient displayed the same pattern but in fewer cells (e.g., CDR2-mutated cells: 28% patient vs. 70% control; **Fig. 3E; Fig. S4A; SuppStat 7**). Among the patient cells with mutations, regional mutation frequencies were ∼50% of control, except for CDR3 with ∼75% (**SuppStat 8–9**). This demonstrates that the observed mutational load in the patient was not dictated by a junctional diversity of CDR3 alone, but involved SHM, though with a substantially lower efficiency and in a smaller subset of cells.

Analysis of clonal architecture reinforced this finding. To quantify repertoire diversity, we applied Hill diversity profiling, which captures both the richness (total number of distinct clones) and the evenness (uniformity of clone sizes) of the B-cell repertoire within a single framework. The patient repertoire exhibited greater clonal diversity and more uniform clone-size distributions across all diversity orders examined (q = 0/1/2) (**Fig. 4A; Fig. S4B; SuppStat 10–11**), indicating a repertoire composed of many similarly sized clones rather than one dominated by large clonal expansions. Consistent with this, the patient harbored fewer expanded clones than the control (8 patient vs. 39 control), and those that were detected were smaller in size and carried a lower *IGH* mutational burden (**Fig. 4B; Fig. S4C-D; SuppStat 12–15**). Lorenz curve analysis revealed a near-uniform clone-size distribution in the patient (Gini = 0.017) compared with the control (Gini = 0.041) (**Fig. S4C**). Phylogenetic reconstruction of individual B-cell clones demonstrated that intraclonal diversification was present in the patient, although markedly less than in the control (**Fig. S4E**). Network analysis of shared CDR3 3-mer motifs revealed convergent CDR3 features across the patient and control, but motif communities were depleted and clone sizes smaller in patient B cells (**Fig. S4F**).

**Figure 4.**
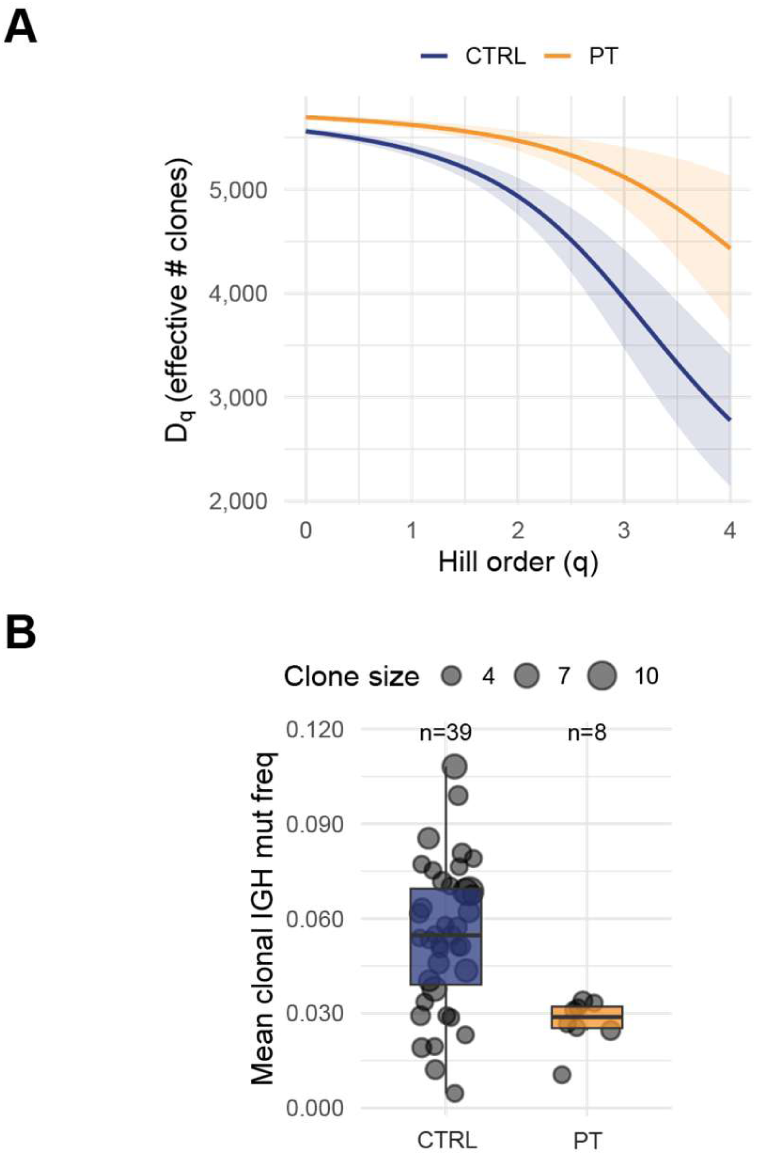
Patient repertoire was more diverse and less dominated; expanded clones were fewer/smaller and less mutated. **(A)** Diversity profiles across Hill orders q (effective number of clones). Patient (PT) clones exceeded control (CTRL) clones in richness (q=0), Shannon (q=1), and Simpson (q=2) diversity with greater evenness and lower dominance (bootstrap tests; p<0.001; Gini and top-clone fractions shown; SuppStat 10–12). **(B)** Expanded clones (≥3 cells/clone) plotted by clone size versus mean IGH mutation frequency. PT contained fewer (8 vs 39) and smaller clones and trended toward lower mean SHM; size–SHM slopes were not significant (CTRL p=0.14; PT p=0.69; SuppStat 13). CTRL: control; PT: patient.

In summary, the patient retains SHM capability but with ∼10-fold lower mutation load, fewer mutated cells, reduced affinity-driven clonal expansion, and altered clonal networks, demonstrating that the p.G197* variant limits SHM efficiency *in vivo*.

### Functional restoration of CSR in patient B cells

To test whether the AID p.G197* variant causes a B-cell-intrinsic defect, we purified naïve B cells from the patient and healthy controls using negative selection and maintained them in iGB cultures **[53]** (**Fig. 5A; Fig. S7A**). In this system, primary human B cells are stimulated on feeder cells expressing membrane-bound CD40L and supplied with IL-21, BAFF, and IL-2. A pulse of IL-4 is additionally added at seeding. Supernatants were collected during culture for antibody measurements, and cells were reseeded weekly to maintain equal starting cell numbers (**Fig. 5B**). Naïve B cells in the iGB cultures undergo staged differentiation: they acquire a GC-like phenotype by ∼day 7, when AID expression peaks **[53, 54]**, and then progress toward a plasmablast-like state by ∼day 14 as AID expression declines.

**Figure 5.**
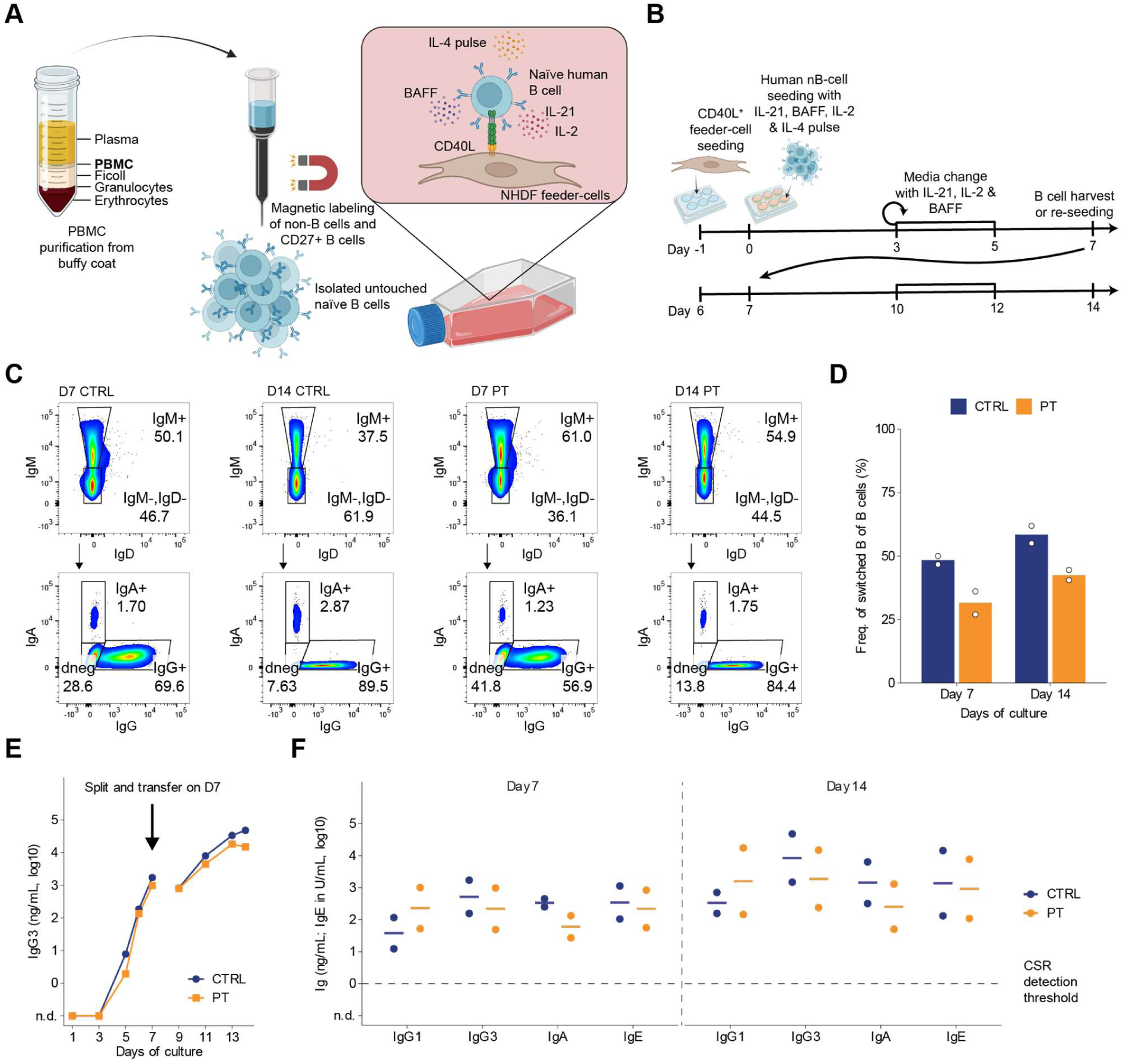
iGB co-culture drives CSR of patient naïve B cells. **(A)** Schematic of the inducible germinal center B-cell (iGB) co-culture: untouched naïve B cells (nB) from the patient (PT; AICDA c.589G>T, p.G197*) and healthy donors (CTRL) were purified by negative selection and seeded on CD40L-expressing NHDF feeder cells with IL-21, BAFF, and IL-2; an IL-4 pulse at seeding biased CSR toward IgG1/IgE. **(B)** Timeline for iGB cultures: cells are harvested on day 7 for reseeding onto fresh feeder cells and maintained to day 14; supernatants collected throughout. **(C)** Flow cytometry of surface immunoglobulin (BCR) isotypes shows CSR in PT B cells under iGB conditions (gating tree in fig S5A), validated with n = 3 different (time-spaced) blood samples (panel and antibodies in supplementary table 1). **(D)** Frequency from (C) of IgM^-^IgD-switched B cells of B cells **(E)** Representative TRIFMA of culture supernatant for IgG3, see more in Fig S6. **(F)** as (E) showing supernatant Ig levels at day 7 and 14 of iGB culture, respectively for IgG1, IgG3, IgA, and IgE. (Coat and detection antibodies can be seen in supplementary table 2). CTRL: control; PT: patient.

In the iGB cultures with strong, continuous activation from CD40L and cytokines, patient naïve B cells proliferated, differentiated and, surprisingly, class-switched their BCRs (**Fig. 5C-D**; **Fig. S5A**). Switching was nonetheless a little less efficient than in control cultures, with switched cells reaching 65% of control frequencies at day 7 and 73% at day 14. Notably, when CD19 positively selected pan B cells were maintained in iGB cultures, patient cells still underwent CSR, albeit delayed and attenuated compared to controls (**Fig. S5B-C**). This delayed CSR for patient pan B cells compared to naïve B cells within iGB cultures, could be explained by competitive pressure from the patient’s unswitched non-naïve cells, dominating the cultures initially.

The isotype-switching was verified for secreted antibodies, where time-resolved immunofluorometric assay (TRIFMA) measurements showed comparable IgG1, IgG3, IgA and IgE secretion within patient cultures compared to controls (**Fig. 5E-F; Fig. S6A-B**). All four switched isotypes were detectable by day 7, and antibody concentrations increased during the second week following reseeding onto fresh feeder cells, demonstrating sustained antibody production by patient B cells.

Collectively, these results show that despite carrying the p.G197* truncation, patient B cells can undergo CSR under strong CD40L- and cytokine-driven activation, indicating that the defect is not absolute and can be functionally overcome in a highly stimulatory microenvironment.

### Reduced AID protein despite normal transcript levels and localization

To understand the allelic contribution of the p.G197* variant in B cells, total *AICDA* transcription and allelic usage were quantified by droplet digital PCR (ddPCR) on control and patient B cell cDNA. Total *AICDA* transcript levels, normalized to *TBP* or *HPRT1*, showed a clear induction that peaked at days 5–7, with similar timing and amplitude in patient and control cultures (**Fig. 6A; Fig. S7B**). Allele-discriminating duplex ddPCR (WT vs variant c.589G>T) demonstrated co-dominant expression in the patient, with the variant comprising 53% of total *AICDA* transcripts by day 7 (**Fig. 6B; Figs. S7C–D**). These data indicate balanced transcription from both *AICDA* alleles and show that the variant transcript is not subject to nonsense-mediated decay.

**Figure 6.**
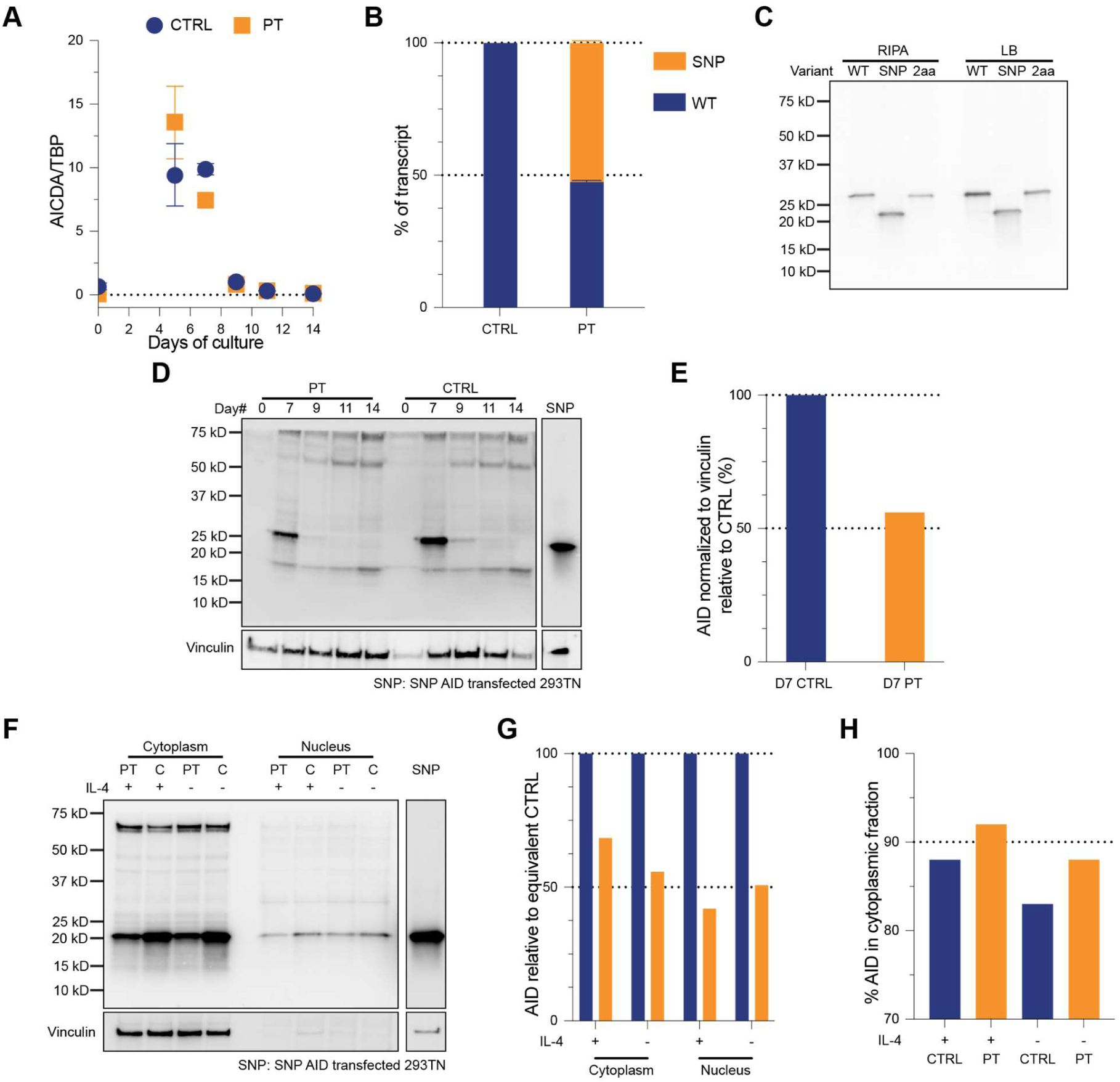
*AICDA* induction is intact for patient B cells, whereas AID abundance is ∼56% of control with maintained cytoplasmic localization. **(A)** ddPCR quantification of total *AICDA* transcription (normalized to TBP) during naïve B-cell iGB culture (days 0, 5, 7, 9, 11, 14), shows similar induction kinetics and amplitude in patient (PT) and control (CTRL) B cells, peaking at days 5–7. (B) Allele-discriminating duplex ddPCR. WT vs c.589G>T (SNP) at day 7 demonstrates codominant expression in PT, with the variant comprising ∼53% of total AICDA transcripts. (C) Validation of antibody AID detectability with HEK293TN transfectants expressing WT, c.589G>T (G197*), or a 2-aa C-terminal truncation. Immunoblots with anti-AID clone mAID-2 detect all constructs under RIPA and load-buffer lysis. WT: Wild type AID, SNP: G197* AID, 2aa: artificial 2 aa C-terminal truncation of AID. Due to Myc-DDK, WT and 2aa have C-terminal FLAG-tag increasing their size. (Antibodies in supplementary table 3). (D) Time-course immunoblot of total AID in primary B cells from PT and CTRL during naïve B-cell iGB culture (days 0, 5, 7, 9, 11, 14). Protein recovered from TRI-reagent preps, probed with anti-AID clone mAID-2; Vinculin as loading control. G197* AID-positive 293TN lysate included as reference. Shows peak protein at day 7. (Antibodies in supplementary table 3). **(E)** Quantification of AID from day 7 cultured naïve B cells (normalized to Vinculin). PT is ∼56% of CTRL abundance. **(F–G)** Subcellular distribution of AID in day 7 iGB B cells (CTRL, PT; ±IL-4). ±IL-4 indicates whether cultures were supplemented with IL-4 as a pulse (-) or continuously (+). Cytoplasmic and nuclear fractions (NE-PER) were immunoblotted for AID (∼24 kDa); Vinculin provided loading control. G197* AID-positive 293TN lysate included. (Antibodies in supplementary table 3). **(G)** Quantification of PT AID-signal, relative to equivalent condition CTRL, from F. Cytoplasmic fractions normalized to Vinculin, nuclear fractions used raw band intensities. AID levels were reduced in PT both cytoplasmic and nuclear extracts, mirroring whole-cell lysates. **(H)** Cytoplasmic/nuclear quantification ratios: Cytoplasmic AID accounts for ∼83–92% of total AID across conditions, consistent with predominant cytoplasmic localization of functional AID. CTRL: control; PT: patient.

Before quantifying AID in patient samples, we verified that monoclonal anti-AID antibodies recognized both wild-type and truncated AID. Using an AID expression plasmid as template and the site-directed mutagenesis technique, two AID variants were generated: the patient’s SNP variant (c.589G>T; p.G197*) and a synthetic two–amino acid (2aa) C-terminal truncation. The WT and 2aa constructs carried a C-terminal FLAG tag, whereas the SNP variant’s premature stop codon prevented FLAG tag inclusion. HEK293TN cells transfected with each construct were analyzed by SDS–PAGE and Western blotting. Both anti-AID clone mAID-2 and clone EK2-5G detected all three proteins: WT, truncated, and patient variant, confirming that their epitopes lie upstream of the two-amino acid C-terminal truncation (**Fig. 6C**, **Fig. S7E**). This validated their use for detecting AID in patient samples.

In primary B-cell iGB cultures, AID protein expression peaked at day 7, paralleling the transcriptional profile observed by ddPCR (**Fig. 6D**). When normalized to vinculin, patient B cells expressed ∼56% of control AID protein levels despite balanced allelic transcript expression (**Fig. 6E**), suggesting altered translation or protein turnover. To assess subcellular distribution, day 7 B cells from patient and control cultures were fractionated into cytoplasmic and nuclear extracts for immunoblotting (**Fig. 6F**). Under both pulse (-) and continuous (+) IL-4 presence, patient AID levels were roughly half of control (42-68%), consistent with the whole-cell data (**Fig. 6G**). Within each condition, cytoplasmic-to-nuclear ratios were comparable between patient and control B cells, with 83–92% of AID localized in the cytoplasm, matching the expected predominantly cytoplasmic distribution (∼90%) **[34, 55]** (**Fig. 6H**).

Together, these data show a normal co-dominant *AICDA* transcription, yet the overall AID abundance is reduced to half without altered subcellular localization.

### Only heterozygous AICDA variants can be rescued in healthy donor cells

To enable investigation of *AICDA* variants in healthy donor B cells, we established targeted disruption of *AICDA* by first optimizing CRISPR/Cas9 guide design and editing conditions. Four gRNAs were evaluated, two in exon 2, for complete AID disruption, and two near the p.G197* variant in exon 5, encoding the AID C-terminal. In SU-DHL-5 and THP-1 cells, both exon 2 gRNAs and the exon 5 gRNA cutting between p.L183/Y184 achieved >95% editing, whereas the proximal p.T195 gRNA showed no detectable activity (**Fig. S8A**). Nucleofection conditions were optimized in primary naïve B cells by delivery of eGFP mRNA across multiple electroporation programs and cell densities, with the optimal condition achieving 78% transfection efficiency and 60% cell viability (**Fig. S8B**). Using this protocol, Cas9 ribonucleoprotein (RNP) delivery yielded >65% editing at the AAVS1 safe-harbor locus following a 3-day pre-activation period (**Fig. S8C**), essential for genome editing in primary B cells **[56]**.

Naïve B cells from two healthy donors were seeded onto feeder cells supplemented with cytokines and nucleofected on day 3 with Cas9 RNPs targeting *AICDA*, achieving 59-88% editing across both donors and all gRNAs (**Fig. 7A, S8D**). Following nucleofection, cells were seeded by limiting dilution into 96-well plate iGB cultures, manually inspected for clonal outgrowth, and expanded into 48-well plates at day 10. Supernatants were collected at day 19 and screened for Ig isotypes by TRIFMA, and selected clones were harvested for genotyping by deep sequencing at day 22. Disruption of exon 2 (Ex2down and Ex2up) severely impaired CSR, with median IgG concentrations falling below the detection limit (1 ng/mL), indicating that the majority of edited clones were unable to undergo CSR (p = 6.7×10^-15^ and p = 1.2×10^-14^, respectively; **Fig. 7B, S8E**). Perturbation of exon 5 (Ex5 Y184) yielded an intermediate phenotype, with median IgG levels remaining above the detection threshold, but a marked reduction in the proportion of CSR-competent clones (p = 5.8×10^-12^; **Fig. 7B, S8E**).

**Figure 7.**
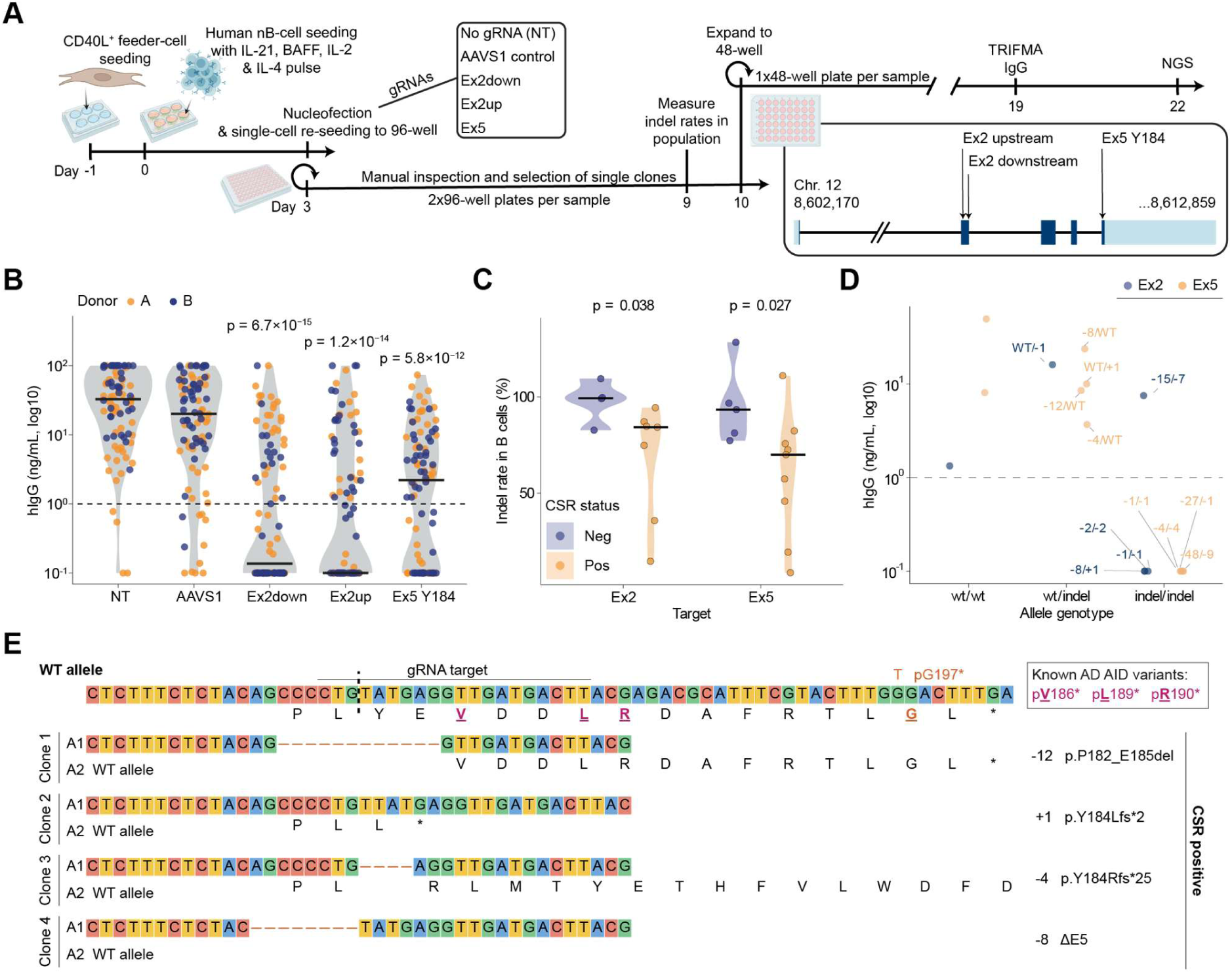
CRISPR/Cas9-mediated disruption of *AICDA* in primary human B cells reveals reduced AID abundance. (**A**) Schematic of the editing and single-cell expansion workflow. Naïve B cells from two healthy donors were pre-activated for 3 days in iGB culture, then nucleofected with Cas9 RNPs targeting AAVS1 (safe-harbor control), *AICDA* exon 2 (Ex2down, Ex2up), or *AICDA* exon 5 (Ex5 Y184). A no-gRNA condition (NT) served as an additional control. Following nucleofection, cells were seeded by limiting dilution into 96-well plates (two plates per condition), manually inspected for clonal expansion, and transferred to 48-well plates at day 10. Supernatants were collected at day 19 for pan-IgG TRIFMA, and selected clones were harvested at day 22 for targeted deep sequencing. (**B**) IgG secretion (log_10_ scale) by individual clonal expansions across editing conditions. Values below 0.1 ng/mL and above 100 ng/mL were capped at these thresholds. Dashed line indicates the assay detection limit (1 ng/mL). Black bars denote medians. Each dot represents an individual clone. Statistical comparisons to the AAVS1 control were performed using van Elteren tests (Wilcoxon rank-sum stratified by donor) with Bonferroni correction for three comparisons. P-values shown are adjusted. (**C**) Indel frequency in genotyped clones stratified by CSR status (Neg, IgG-negative; Pos, IgG-positive) for exon 2 and exon 5 targets. Black bars denote medians. Statistical comparison between CSR-negative and CSR-positive clones were performed using two-sided Welch’s t-tests. (**D**) IgG concentration of individual exon 2-and exon 5-edited clones stratified by genotype. Allele categories were assigned based on deep sequencing read ratios: wild-type homozygous (wt/wt), heterozygous (wt/indel), biallelic (indel/indel), or multiclonal (multi). Dashed line indicates the detection limit. (**E**) Allele-level characterization of four CSR-competent, heterozygous exon 5-edited clones. Each carries one wild-type allele and one indel allele: Clone 1, 12-bp in-frame deletion (p.P182_E185del); Clone 2, 1-bp insertion (p.Y184Lfs2); Clone 3, 4-bp deletion (p.Y184Rfs25); Clone 4, 8-bp deletion predicted to cause exon-5 skipping (ΔE5).

Clones exhibiting strong expansion, proliferating to 5-15×10³ cells were selected for genotyping by PCR amplification and deep sequencing. To assess genotyping reproducibility, a subset of eight clones was re-amplified and sequenced independently, yielding highly concordant results (Pearson r = 0.996; **Fig. S8F**). CSR-negative clones carried significantly higher indel frequencies than CSR-positive clones for both exon 2 (p = 0.038) and exon 5 (p = 0.027) targets, with CSR-negative clones uniformly showing >76% indels and CSR-positive clones displaying a broad range (9-111%), where values exceeding 100% reflect biallelic editing in B cells detected by deep sequencing of alleles from both B cells and feeder cells (**Fig. 7C**). Allele-level analysis revealed that clones harboring at least one wild-type allele (wt/wt or wt/indel) retained the capacity for IgG secretion, whereas indel/indel clones generally lost CSR (**Fig. 7D**). Notably, an exon 2 indel/indel clone carrying an in-frame 15-bp deletion retained CSR, suggesting the deletion preserved sufficient AID activity. By contrast, an exon 5 indel/indel clone with in-frame 9-bp and 48-bp deletions remained CSR-negative, indicating that frame preservation alone is insufficient when the deleted residues lie within the C-terminal region required for CSR. Among CSR-competent exon 5-edited clones, the indel allele varied, including an exon-5-skipping deletion (ΔE5), frameshift truncations (e.g. p.Y184Lfs2, p.Y184Rfs25), and an in-frame deletion (p.P182_E185del), while at least one wild-type allele was preserved (**Fig. 7E, S9A**).

Together, these data show that CSR was largely undetectable in clones carrying biallelic out-of-frame indels, whereas clones retaining one wildtype allele remained CSR-competent irrespective of whether the edited allele targeted exon 2 or the C-terminal-encoding exon 5.

## Discussion

We identify AICDA c.589G>T (p.G197*) as a minimal C-terminal truncation of AID that abolishes CSR and markedly reduces SHM. This variant drives adult-onset HIGM2 in a 34-year-old female presenting with recurrent respiratory infections and unswitched memory B cells. Both her adult onset and her asymptomatic heterozygous father align with dominant rather than recessive AID deficiency, since AR disease typically presents in early childhood **[14]**, whereas AD C-terminal variants frequently present in adulthood and segregate within families alongside mildly affected carriers **[19, 20, 38]**.

Single-cell repertoire sequencing revealed a broad but weakly expanded clonal landscape, with MBCs carrying few or no V-region mutations across CDR2, FWR3, and CDR3. When mutations were present, levels were reduced to ∼50–75% of controls. The reduced frequency of mutated cells, small clone sizes, and diminished intraclonal diversification indicate defective initiation or propagation of SHM rather than an inability of AID to catalyze deamination once engaged. Thus, SHM remains operative but inefficient in the patient’s B cells carrying p.G197*.

Although both AICDA alleles were expressed at similar levels, total AID protein abundance was reduced to ∼56% of control levels, suggesting altered translation or protein turnover. The C-terminus contributes to nuclear export and eEF1A-dependent cytoplasmic retention, and nuclear AID undergoes rapid proteasomal degradation **[33, 57, 58]**. Impaired nuclear export or cytoplasmic retention could therefore promote increased degradation of truncated AID in the nucleus, which would explain the reduced abundance. A p.G197A substitution has been reported to likewise impair CSR while retaining a predominantly cytoplasmic distribution, illustrating that C-terminal alterations can impair AID function without a detectable change in subcellular localization **[24].**

Reduced abundance alone is unlikely to explain the CSR defect. Heterozygous carriers of recessive AICDA alleles, including deletion of an entire AICDA coding allele, can maintain normal serum immunoglobulins and intact *in vitro* CSR **[38]**, although SHM can be reduced in their IgG MBCs **[39]**. These observations argue against simple haploinsufficiency as a sufficient explanation and favor an additional effect of the truncated protein. At the same time, the low abundance of mutant AID places strong constraints on which dominant-negative mechanisms remain possible. Models requiring abundant mutant protein such as inactive heterodimerization with wild-type AID **[22]**, sequestration of shared cofactors **[35]**, or formation of nuclear condensates that withdraw wild-type AID from the IGH locus **[36]** are all inconsistent with the levels we measure. Each of these mechanisms requires the mutant to be present at levels at least matching wild-type AID, and co-condensation specifically requires a high nuclear mutant-to-wild-type ratio **[36]**. The measured total AID level corresponds to a single allele, which makes such an excess unlikely.

Studies of larger C-terminal deletions suggest an alternative. Truncated AID can retain deaminase activity while impairing the protection and repair of S-region DSBs, and its dominant-negative activity can depend on catalysis rather than on abundance **[26, 37]**. Two genetic controls support this distinction. Catalytically inactive full-length AID does not interfere with CSR even when expressed above the level of the truncated protein, and inactivating the catalytic site of a C-terminally truncated AID abolishes its dominant-negative effect while leaving single-stranded DNA binding intact, arguing against both molar excess and substrate competition **[26]**. Mechanistically, breaks generated by truncated AID show reduced recruitment of end-protection factors at Sμ alongside increased association of resection-associated factors such as RPA. Because unprotected ends are degraded into long single-stranded tracts that non-homologous end joining cannot ligate, these breaks are instead resolved by homologous recombination or other rearrangements that compete with productive CSR. Limiting DNA end resection partially restores CSR, supporting a role for excessive resection in the switching defect **[26]**. The dominant-negative effect is also lost in Msh2-deficient B cells, suggesting that it requires MSH2-dependent processing of AID-induced lesions into DSBs **[37]**. We therefore propose that p.G197* may combine reduced effective AID availability with impaired coupling of deamination to CSR productive repair, such that transient mutant molecules deaminate DNA during brief nuclear presence but fail to recruit the downstream end-protection and repair complexes required to resolve switch-region DSBs. Under this model, lesions generated by truncated AID could interfere with CSR despite the presence of wild-type AID.

Remarkably, the recovery of CSR in iGB cultures suggests that any such interference can be partly overcome under sustained B cell activation. Class-switched B cells accounted for at least 21% of patient B cells at day 7 and 38% at day 14, and IgG1, IgG3, IgA and IgE secretion was comparable to controls. Recovery occurred despite reduced total AID abundance, indicating that normalization of protein levels was not required for CSR under these conditions. CRISPR-Cas9 editing of healthy donor B cells provided complementary evidence. Clones retaining a wild-type AICDA allele remained CSR-competent despite exon-5 editing, whereas clones with biallelic out-of-frame indels generally lost switching, indicating that one intact allele can support CSR in the iGB system despite C-terminal disruption of the other. The condition that differs most from an *in vivo* germinal center is instead the absence of competition and affinity-based selection. Cells that have sustained AID-dependent damage are supported by continuous CD40L and survival cytokines rather than eliminated, so unproductive DSBs that would be counter-selected *in vivo* survives. Sustained stimulation may additionally compensate for inefficient CSR by maintaining activation and extending the time available for productive recombination, and differences in stimulation, feeder-cell support and culture duration may likewise explain the greater recovery observed here than in previous studies of C-terminal variants **[17, 38]**. Consistent with a selection-dependent component, CSR was delayed and attenuated when the patient’s pan B cell compartment was cultured in place of purified naïve B cells, and the incomplete recovery may represent the residual dominant-negative effect persisting under maximal activation.

The clinically unaffected carrier father suggests incomplete penetrance, consistent with the variable expression reported for other C-terminal *AICDA* variants **[20, 38]**. Although trio WGS did not identify additional variants relevant to humoral immunity, polygenic background and environmental factors may influence disease expression. The adult presentation also underscores the relevance of *AICDA*-associated disease in patients without an early history of immunodeficiency. Clinically, management should emphasize optimized immunoglobulin replacement and infection prophylaxis; allogeneic hematopoietic stem-cell transplantation (HSCT) remains reserved for clear indications and is best established for CD40L/CD40 deficiency **[59, 60]**.

Our findings associate loss of the final two residues of AID with profound impairment of CSR and markedly reduced SHM. Reduced AID abundance may contribute alongside dominant-negative interference. The substantial recovery of CSR *ex vivo* highlights retained functional capacity and identifies activation conditions as an important determinant of the phenotype.

## Methods

### Bone marrow examination

Bone marrow evaluation included Jamshidi core biopsy, aspirate smears, imprint preparations, and clot section. Histological and cytological analyses were performed using standard staining and immunohistochemical methods, including MUM1 immunostaining and *in situ* hybridization for plasma cell clonality.

### Whole Exome Sequencing (WES) and Whole Genome Sequencing (WGS)

Genomic DNA was extracted from EDTA-anticoagulated peripheral blood using the QIAsymphony DSP DNA Midi Kit (Qiagen). WES libraries were prepared with the TruSeq DNA Sample Preparation Kit (Illumina) and enriched with the EZ Human Exome Library v3.0 (Roche NimbleGen). WGS libraries were generated using the DNA PCR-Free Prep, Tagmentation kit (Illumina). All libraries were sequenced with paired-end 150 bp reads and dual indexing (WES on Illumina NextSeq 500, WGS on Illumina NovaSeq 6000). Sample identity was confirmed using a SNP genotyping panel.

Quality thresholds were >95% of targeted bases at ≥30× coverage for WES (median >80×), and >95% of the genome at ≥10× coverage for WGS (median >30×). Reads were aligned to the human reference genome (WES: GRCh37/hg19; WGS: GRCh38/hg38), and small variants were called with GATK HaplotypeCaller (Broad Institute). Structural variants were assessed using ExomeDepth (WES) or Manta, CNVnator, Lumpy, and Delly (WGS).

WES variants were analyzed as a singleton using Ingenuity Variant Analysis (Qiagen), applying a curated panel for antibody deficiency genes (adapted from Tangye et al.) **[61]** and allele frequency cutoffs of <1% for homozygous and <0.01% for heterozygous variants in gnomAD. WGS variants were evaluated as a trio using VarSeq (Golden Helix), filtering for candidate variants consistent with autosomal dominant (*de novo*) and autosomal recessive (homozygous or compound heterozygous) inheritance, with the same allele frequency thresholds.

### Purification of PBMCs and primary human naïve and pan B cells

Peripheral blood mononuclear cells (PBMCs) were isolated from whole blood or buffy coats obtained from the patient or anonymized healthy donors (Aarhus University Hospital Blood Bank, Denmark). Samples were diluted 1:1 with PBS and layered over Ficoll-Paque PLUS (GE Healthcare) in SepMate tubes (STEMCELL Technologies), then centrifuged at 1,200 g for 10 min at 20 °C. The PBMC layer was collected, washed once in PBS (500 g, 10 min, 20 °C), incubated in PBS containing 27 mM EDTA for 5 min at room temperature, centrifuged (300 g, 8 min, 20 °C) and washed again in PBS. Cells were resuspended in 5 mL MACS buffer (PBS, 2% heat-inactivated FBS, 2 mM EDTA) at room temperature, counted, and approximately 4 × 10^8^ cells were reserved for magnetic separation.

For naïve B-cell isolation, PBMCs were first blocked with Human BD Fc Block (clone Fc1; BD 564220) for 5 min on ice, then incubated with the Miltenyi Biotec Human Naïve B Cell Isolation biotin-Ab cocktail (130-091-150) for 30 min. Cells were diluted with MACS buffer and centrifuged (200 g, 10 min, 4 °C), the pellet was resuspended in 1 mL MACS buffer, and anti-biotin MicroBeads (Miltenyi Biotec) pre-diluted in 2 mL MACS buffer were added for 20 min. The suspension was passed through a 70 µm cell strainer onto pre-wetted LS columns (Miltenyi Biotec), washed twice with 3 mL MACS buffer, and the flow-through was collected as the untouched naïve B-cell fraction. Phenotype (CD19+, CD27−, IgD+) was confirmed by flow cytometry with a minimum purity of 98%.

For pan B-cell (CD19^+^) purification used in supplementary cultures and single-cell sequencing, PBMCs were processed as above but with positive selection using Miltenyi Biotec Human CD19 MicroBeads (130-050-301) according to the manufacturer’s instructions; the eluate (positive fraction) was collected. Purity was assessed by flow cytometry (CD19+, CD27±, IgD±) with a minimum purity of 98%. Aliquots of PBMCs as well as both naïve and pan B cells were cryopreserved in 90% FBS and 10% DMSO using Corning CoolCell containers following the manufacturer’s recommendations.

### Flow cytometry

Cell suspensions (PBMCs, freshly MACS-purified human B cells for purification validation, or B cells harvested from iGB cultures for CSR analysis) were prepared in flow cytometry (FC) buffer (PBS, 2% FBS, 1 mM EDTA). Compensation matrices were generated using anti-mouse Igκ particles (BD, 552843). For staining, 100 µL of sample was pre-incubated with 20 µL Fc block (5–10 min, on ice), followed by surface staining with 100 µL of appropriate fluorophore-conjugated Ab cocktail (**Supplementary Table 2**) prepared 1:1 in FC buffer and Brilliant Stain Buffer (BD, 563794). After 30 min on ice, cells were washed (200 g, 5 min, 4 °C) and resuspended in FC buffer.

For intracellular detection of Ig isotypes (CSR panel), additional staining was performed using the eBioscience Fixation/Permeabilization Buffer Set (#00-5523-00) according to the manufacturer’s instructions. Briefly, cells were fixed in formalin fixation buffer for 30–60 min at room temperature, centrifuged (350 g, 5 min, 21 °C), washed once with permeabilization buffer (350 g, 5 min, 21 °C), and resuspended in 100 µL permeabilization buffer containing 2% FBS. A 100 µL working solution of intracellular Abs was added and incubated for 30 min at room temperature. Cells were then washed once with permeabilization buffer (350 g, 5 min, 21 °C) and returned to FC buffer.

Prior to acquisition, all samples were passed through a 100 µm cell strainer to remove aggregates. Data were acquired on a NovoCyte Quanteon 4025 (Agilent) equipped with four lasers (405, 488, 561, and 640 nm) and 25 detectors and analyzed in FlowJo v10.10.0 (BD).

### In silico analyses of AICDA variants

Variants previously reported in patients with HIGM2 were retrived using PubMind **[40]**. Structural modelling of AID was performed using AlphaFold2 (AlphaFold Monomer v2.0), retrieving the predicted 3D model (AF-Q9GZX7-F1-v6) and per-residue predicted Local Distance Difference Test (pLDDT) scores **[62]**. AlphaMissense variant-effect predictions for all possible missense substitutions in AID were obtained from the AlphaMissense database using the UniProt identifier Q9GZX7 **[50, 63]**. ProteoCast was used to extract GEMME evolutionary constraint scores and segment boundaries for each residue **[64, 65]**. Amino acid sequences of AID orthologs from representative vertebrates: human (*Homo sapiens*), rhesus macaque (*Macaca mulatta*), mouse (*Mus musculus*), chicken (*Gallus gallus*), Chinese softshell turtle (*Pelodiscus sinensis*), African clawed frog (*Xenopus laevis*), and Japanese pufferfish (*Takifugu rubripes*) were downloaded from UniProt and aligned using Clustal Omega **[66]**. Human genetic variation data were obtained from gnomAD v4.1.0, and pathogenic or likely pathogenic variants were taken from ClinVar (downloaded November 3rd, 2025). ClinVar variants were further annotated with CADD scores using the CADD v1.6 scoring interface **[67]**.

### Single-cell transcriptomic evaluation of CSR and SHM

Freshly purified CD19+ B cells from CTRL and PT were processed side-by-side with the 10x Genomics Chromium next GEM Single Cell 5′ Reagent Kit v2 (PN-1000263) for gene expression and the Chromium Single Cell Human BCR Amplification Kit (PN-1000253) for BCR sequencing. Libraries were prepared per manufacturer instructions and sequenced on an Illumina NovaSeq (target ∼30,000 reads per cell). FASTQs were processed with Cell Ranger 8.1.0 to generate gene-barcode matrices with genome reference GRCh38-2024-A, and V(D)J contigs with V(D)J reference vdj_GRCh38_alts_ensembl-7.1.0.

Gene-barcode matrices were analyzed using Seurat v5.3.0 **[51]**. Cells were filtered on gene/UMI counts and mitochondrial fraction, and doublets were removed using DoubletFinder **[68]**. Counts were log-normalized, 3,000 highly variable genes were identified, and IGHV/IGKV/IGLV gene segment usage was excluded from scaling. PCA and SNN clustering were performed, with clustree **[69]** guiding resolution choice.

Downstream analyses were restricted to B-lineage clusters, with CTRL and PT jointly analyzed after anchor-based CCA integration, followed by UMAP visualization. Differential expression was performed with Wilcoxon tests and BH correction.

For BCR analysis, V(D)J contigs were assigned with IgBLAST v1.22.0 against the IMGT human Ig germline reference (downloaded March 2025), then processed with the Immcantation suite **[52]** for AIRR schema conversion, clonal assignment (SCOPer), germline reconstruction, SHM quantification (SHazaM/Alakazam), and lineage tree inference (Dowser, visualized with ggtree). Analyses were restricted to cells with one productive IGH paired to a single IGK or IGL. Per-cell BCR annotations, including isotype (c_call) and heavy-chain mutation frequency, were merged into the Seurat object, enabling integrated visualization and state-specific SHM comparisons (nB, MBC, PC). Region-level SHM summaries (IMGT FWR/CDR segments) were computed for group comparisons. Statistical procedures are detailed under Statistics.

### Human primary B-cell cultures

CD19^+^ pan and naïve B cell cultures with CD40L-expressing NHDF feeder cells were performed as in Rovsing & Green *et al.* 2025 **[53]**, with the addition of IL-2. Feeder cells were expanded at least 8 days in RPMI-1640 [+] L-Glutamine with 1% P/S (invitrogen), and 10% FBS (Sigma), then seeded the day before B-cell addition and held overnight in B-cell medium (BCM: RPMI-1640[+] L-Glutamine + 1% P/S, 10 mM HEPES, 1 mM sodium pyruvate, 55 µM 2-mercaptoethanol (Invitrogen), 10% FBS). On day 0, human B cells were seeded onto the feeder cells in BCM supplemented with final 10 ng/mL IL-4, IL-21, BAFF (Peprotech, 200-04, 200-21 and 310-13, respectively), as well as 50 ng/mL IL-2 (Peprotech 200-02). On day 3 and 5, half of the medium was replaced with fresh BCM similarly supplemented with IL-21, BAFF and IL-2. For subcellular localization, the continuous (+) IL-4 conditions additionally included IL-4 in media change. On day 7, B cells were harvested or re-seeded onto fresh feeder cells and maintained for an additional week under the same regimen. Cultures were kept at 37 °C, 5% CO₂. Supernatants collected at indicated time points were stored with 0.1% (w/v) sodium azide for TRIFMA isotype quantification.

### TRIFMA

To quantify secreted class-switched Abs within either serum samples or culture supernatants, time-resolved immunofluorometric assays (TRIFMAs) were performed, with all values determined in triplicate. FluoroNunc MaxiSorp 96-well plates were coated overnight 4 ℃ with 100 µL/well of capture Ab in PBS: 0.3 µg/mL donkey anti-human IgG (Jackson ImmunoResearch, 709-005-098), 2.5 µg/mL mouse anti-human IgG1 (SouthernBiotech, 9052-01), 1 µg/mL mouse anti-human IgG3 (SouthernBiotech, 9210-01), 1 µg/mL goat anti-human IgA (SouthernBiotech, 2053-01), or 2.5 µg/mL mouse anti-human IgE (SouthernBiotech, 9240-01).

Plates were blocked for 1 h at room temperature with 200 µL/well TBS (Tris-buffered saline: 137 mM NaCl, 2.7 mM KCl, 25 mM Tris/Tris-HCl) containing 0.09% (w/v) sodium azide (Fisher Scientific, BP2471-500) and 1% BSA (Sigma A 4503), then washed 3x using TBS with 0.05% Tween-20 (TBS-TW). Samples were diluted 1/5 in TBS-TW/0.1% BSA and added in duplicates for 2 h incubation at RT (1 h for pan-IgG). Standards were prepared in TBS-TW/0.1% BSA diluting either; human IgG (gammanorm, 478393) 2000 ng/mL with seven 5-fold dilutions; human IgG1 (SouthernBiotech, 0151-K) and IgG3 (SouthernBiotech, 0153-L) 100 ng/mL with ten 2-fold dilutions; human IgA (SouthernBiotech, 0155-L) 500 ng/mL with ten 2-fold dilutions; for IgE, human plasma diluted 1/3 and further by ten 2-fold dilutions. After 3x TBS-TW washes, appropriate biotinylated detection Abs were applied at room temperature: goat anti-human IgG (H+L) (SouthernBiotech, 2016-08) at 0.1 µg/mL for 1 h (pan-IgG or IgG1 detection) or 2 h (IgG3); goat anti-human IgA (Invitrogen, A18791) at 0.5 µg/mL for 2 h; mouse anti-human IgE (SouthernBiotech, 9250-08) at 0.2 µg/mL for 2 h. Plates were washed 3x and incubated 30 min at RT with Eu^3+^-labeled streptavidin (PerkinElmer, 1244-360), at 1 µg/mL in TBS-TW with 25 µM EDTA. Lastly, plates were washed 3x, then developed with 200 µL/well Enhancement Solution (Ampliqon). After 5 min shaking, time-resolved fluorescence was measured on Victor X5 plate reader (Perkin Elmer).

### ddPCR

Total RNA was extracted from cultured primary B cells using Direct-zol RNA MicroPrep (Zymo research, R2060) according to the manufacturer’s instructions, including on-column DNase I digestion. RNA was quantified by spectrophotometry. cDNA was generated with the iScript cDNA Synthesis Kit (Bio-Rad) in 20 µL reactions.

Absolute *AICDA* transcript copies were quantified on a QX200 Droplet Digital PCR System (Bio-Rad) using duplex hydrolysis-probe assays with *AICDA* in FAM-channel target and either *TBP* or *HPRT1* as HEX-channel reference.

Per 20 µL well we used 10 µL ddPCR Supermix for Probes (no dUTP), 1 µL 20x *AICDA* assay (FAM), 1 µL 20x reference assay (HEX), 1 µL cDNA, and 7 µL water. Droplets were generated on the QX200 droplet generator, PCR-amplified under standard probe-based conditions (enzyme activation 95 °C 10 min; 40 cycles of 94 °C 30 s and 60 °C 60 s; enzyme deactivation 98 °C 10 min; 4 °C hold; 2 °C/s ramp; heated lid). Droplets were read on the QX200 reader and analyzed in QuantaSoft; wells with ∼12–15k accepted droplets were typical. For these expression runs, auto-thresholds were accepted with visual verification. *AICDA* copies/µL were normalized to the reference *TBP* or *HPRT1*.

Allele-discriminating duplex assays were used to quantify WT vs PT variant transcripts (HEX-WT in Ch2; FAM-Mut in Ch1). 20 µL reactions contained 10 µL ddPCR Supermix for Probes (no dUTP), 1 µL allele-discrimination primer/probe mix (both probes present), 1 µL cDNA, and 8 µL water. High-expressing cDNAs were diluted 1:1 in water before. Droplet generation and cycling matched the expression assay. Because the two positive clouds partially overlap in amplitude space, thresholds for allele calls were set manually in QuantaSoft based on the four-cluster pattern (Ch1+/Ch2−, Ch1−/Ch2+, double-positive, double-negative) with no-RT/NTC control guidance. Mutant fractional abundance was calculated as Mut/(Mut+WT)×100%; for presentation as well as the reciprocal (WT/(WT+Mut)).

### Plasmid construction (AICDA variants)

Human *AICDA* (pCMV6-Entry-AICDA, KanR, Myc-DDK-tagged) was edited by site-directed mutagenesis (QuikChange Lightning, Agilent). For the c.589G>T SNP, primers hAICDA-G589T-F (5′-CGCATTTCGTACTTTGTGACTTACGCGTACGCG-3′) and hAICDA-G589T-R (5′-CGCGTACGCGTAAGT CACAAAGTACGAAATGCG-3′) were used; a two–amino-acid C-terminal deletion was generated with primers AID-deletion-F (5′-AGACGCATTTCGTACTTTGACGCGTACGCGG-3′) and AID-deletion-R (5′-CCGCGTACGCGTCAAAGTACGAAATGCGTCT-3′). Each 50-µL reaction contained 5 µL 10× buffer, 50 ng plasmid template, 125 ng of each primer, 1 µL dNTP mix, 1.5 µL QuikSolution, water to 49 µL, and 1 µL Lightning enzyme. Cycling: 95 °C 2 min; 18 cycles of 95 °C 20 s, 60 °C 10 s, 68 °C 2 min 45 s (5.5-kb plasmid); final extension 68 °C 5 min. Products were treated with DpnI (2 µL, 37 °C, 20 min), transformed into TOP10, selected on kanamycin (25 µg/mL), and miniprepped. Edits were verified by Sanger sequencing using vector primers; two independent colonies per construct were sequence-confirmed.

### HEK293TN AICDA variant transfection

HEK293TN were seeded at 2.5 × 10⁵ cells/well (12-well, 1 mL) one day before transfection. Per well, complexes were made by mixing 8 µL Lipofectamine 2000 in 250 µL Opti-MEM with 2 µg plasmid DNA in 250 µL Opti-MEM (20–30 min at RT) and adding the 500 µL mix to cells. WT *AICDA*, the c.589G>T SNP, and the 2-aa truncation were transfected in parallel. Cells were harvested 24-48 h post-transfection for western blotting.

### Western blotting

Cells were lysed and analyzed by SDS–PAGE followed by semi-dry transfer to nitrocellulose (7 min). Membranes were blocked in TBS with 0.1% Tween-20 (TBS-TW) and probed overnight (4 °C) with anti-AID rat mAb EK2-5G9 (#4959, 1:1000) or biotinylated anti-AID mAID-2 (eBioscience 13-5959-82, 1:1000) prepared in TBS-TW supplemented with HSA (1 mg/mL), normal human Ig (100 µg/mL), and EDTA (1 mM). After TBS-TW washes, EK2-5G9 blots were developed with an HRP-conjugated secondary Ab (1:3000, 2 h, RT); mAID-2 blots used HRP-streptavidin (DAKO P0397, 1:3000, 2 h, RT). Signals were detected by ECL, imaged on an ImageQuant LAS-4000 (GE Healthcare), and quantified in ImageJ (v1.54p) using consistent ROI widths and area-under-curve lane integration, with normalization to loading controls.

For HEK293TN transfectants (WT AID, c.589G>T, and a 2-aa ΔC), cells were harvested 24 h post-transfection, washed once in PBS, and lysed either directly in sample/loading buffer (“LB”) or in RIPA. For gel loading, RIPA lysates were mixed with sample buffer, TBS, and 1 M DTT at 5:2.5:1.5:1 µL and boiled 3 min; LB lysates were mixed 5:4:1 µL (lysate:TBS:DTT) and boiled 3 min. Equal volumes were loaded. Note that the C-terminal FLAG-tag (from Myc-DDK) increases size of WT and 2-aa ΔC AID, while SNP (c.589G>T) truncates it.

For time-course analysis of total AID in primary B-cell iGB cultures (PT and CTRL), ∼0.5 × 10⁵ cells were collected at days 0, 5, 7, 9, 11, and 14 into PBS and TRI-reagent; protein was recovered by acetone precipitation from the RNA workflow. Reduced samples (25 µL/lane) were resolved and transferred as above. Membranes were cut at ∼75 kDa to allow parallel probing: the lower portion with biotinylated (mAID-2) anti-AID (then HRP-streptavidin), and the upper with anti-Vinculin (mouse mAb, Sigma V9131, 1:10 000) followed by HRP anti-mouse (DAKO P0260, 1:3000). A 293TN AID-positive lysate (2 µL; ∼24 kDa band) was included as a control.

For subcellular localization, day 7 iGB-cultured patient or control B cells (±IL-4; ∼1 × 10⁵) were fractionated using NE-PER Nuclear and Cytoplasmic Extraction Reagents (Thermo) per manufacturer’s instructions. Cytoplasmic (∼105 µL) and nuclear (∼50 µL) fractions were reduced, boiled 3 min, and 25 µL loaded. Blotting and Ab development were as above (anti-AID, mAID-2, on the lower membrane; anti-Vinculin on the upper). AID was expected at ∼24 kDa and Vinculin at ∼116 kDa throughout.

### Cell line cultures

SU-DHL-5 and THP-1 cells were maintained in RPMI-1640 [+] L-Glutamine (Lonza) supplemented with 10% FBS (Sigma) and 1% P/S (Invitrogen). SU-DHL-5 and THP-1 cell lines were purchased from the German Collection of Microorganisms and Cell Cultures (DSMZ) and ATCC, respectively.

### Nucleofection of cell lines and human primary B cells

Nucleofection was performed using the Lonza 4D-Nucleofector with 20 µL Nucleocuvette strips. Cells were pelleted at 200 × g for 5 minutes, washed once in the appropriate buffer (Opti-MEM for THP-1 and SU-DHL-5; 1M buffer for primary B-cells (5 mM KCl, 15 mM MgCl₂, 120 mM Na₂HPO₄/NaH₂PO₄, 50 mM mannitol)), and resuspended in 20 µL. For mRNA delivery, cells were mixed with 1 µg *in vitro*–transcribed eGFP mRNA. For CRISPR editing, Cas9 RNP complexes were assembled by incubating 3.2 µg chemically modified gRNA (Synthego) (**Supplementary Table 4**) with 6 µg Cas9 protein (Alt-R SpCas9, IDT) for 15 min at 25°C and then combined with either 2×10^5^ primary B cells or 6×10^5^ THP-1/SU-DHL-5 cells. Nucleofection was carried out using program CM-138 with P3 solution, after which 100 µL pre-warmed medium was added and the full volume transferred to the appropriate culture plate. Cell viability and eGFP expression were assessed after 24 h, and editing efficiency was measured by Sanger sequencing after 5 days.

For primary B-cell editing, naïve B cells were pre-activated for 3 days in iGB co-culture before nucleofection. Cells were then either seeded by limiting dilution onto feeder cells or were returned to bulk iGB culture. Bulk cultures were collected on day 9 for genomic DNA extraction and Sanger sequencing. Single-cell-derived clones were expanded from 96-well plates to 48-well plates on day 10, screened for pan-IgG secretion on day 19, and collected on day 22 for genomic DNA isolation and deep sequencing.

### Editing analysis by Sanger sequencing and deep sequencing

Genomic DNA was obtained by direct lysis of cell pellets in QuickExtract (Biosearch Technologies). Pellets were resuspended at 0.5-1×10^4^ cells per µL, incubated at 60°C for 6 minutes and 100°C for 10 minutes, and the crude lysate was used directly for PCR. Targeted loci were amplified with primers flanking the CRISPR cut site (**Supplementary Table 5**), purified, and subjected to Sanger sequencing. Chromatograms were analyzed using the ICE platform (Synthego) to estimate indel frequencies.

For deep sequencing of expanded primary B-cell clones, genomic DNA was extracted with QuickExtract (Biosearch Technologies) and amplified using primers (**Supplementary Table 6**). PCR was performed with Phusion Plus PCR Master Mix (Thermo Scientific) using 1 µL lysate in 50 µL reactions for 30 cycles. Amplicons were purified with HighPrep PCR Clean-up System (MagBio) beads according to manufacturer’s protocol. DNA concentrations were measured on a Qubit fluorometer (Thermo Scientific). Then, indexed libraries were generated in a second PCR using TruSeq adapters (Phusion Plus, 25 µL reaction, 4 ng template from PCR1, 8 cycles), purified with the HighPrep PCR clean-up system, quantified by Qubit, pooled, and sequenced using 150-bp paired-end reads on an iSeq 100 (Illumina).

All samples achieved at least 5×10^3^ aligned reads. Overlapping paired-end reads were merged using FLASH **[70]** with a minimum overlap of 5 bp and a maximum overlap of 145 bp, yielding consensus reads of approximately 247–248 bp. To distinguish reads originating from primary B-cell clones from those derived from feeder cells, we exploited heterozygous SNPs within the amplicon sequences as discriminating identifiers where available, or sequenced HLA-A amplicons unique to each donor (**Supplementary Table 7**). Specifically, merged reads were genotyped at the discriminating SNP position by identifying a conserved key sequence flanking the SNP and calling the allele based on the immediately adjacent nucleotide. For *AICDA* exon 2, an A/G SNP was used to separate alleles; for *AICDA* exon 5, an A/C SNP was used. Donor A had a discriminating SNP in exon 5 but not exon 2, and donor B had a distinguishing SNP in exon 2 but not exon 5. For samples lacking an intronic discriminating SNP, HLA-A amplicons were genotyped at two independent regions to assign reads to donor or feeder origin. Only samples with more than 50% B-cell-derived compared to feeder cell-derived reads were included in downstream analysis. Editing rates were quantified using CRISPResso2 with standard parameters except that the quantification window was expanded with 3 bp **[71]**.

Where a discriminating SNP was present at the target locus (donor A exon 5, donor B exon 2), per-allele editing rates were estimated independently: the donor-specific allele rate was used directly, while the shared-allele rate was corrected by dividing by the donor-derived read fraction, and the two chromosomal estimates were averaged. Where no discriminating SNP was available at the target locus (donor A exon 2, donor B exon 5), the overall indel rate was divided by the donor-derived read fraction to obtain the B-cell-specific editing rate. Clones were classified based on the donor-frame allele distribution as wild-type homozygous (wt/wt), heterozygous (wt/indel), biallelic (indel/indel), or multiclonal (multi), called homozygous when one allele covered ≥75% of donor reads, heterozygous when the top two combined did, and multi otherwise. A subset of clones was independently re-amplified and sequenced to assess genotyping reproducibility (**Supporting data values**).

### Sex as a biological variable

Sex was not considered as a biological variable in experiments using cell cultures. For scRNAseq sex-chromosomal genes were assessed, and their exclusion did not affect the results.

### Statistics

Statistical analyses were carried out in R (v4.5.1) and GraphPad Prism v10.5.0. In R, we used rstatix for Wilcoxon tests with Benjamini–Hochberg correction, including Hodges–Lehmann effect-size estimation, alongside differential expression testing within Seurat and mutation/clonal analyses within the Immcantation framework **[51, 52, 72, 73].** For the 48-well clonal expansion assay (**Fig. 7B**), hIgG concentrations between editing conditions and the AAVS1 control were compared using van Elteren tests (Wilcoxon rank-sum stratified by donor) with Bonferroni correction for three comparisons, implemented using the coin R package. Indel frequencies in CSR-positive versus CSR-negative clones (**Fig. 7C**) were compared using two-sided Welch’s t-tests; where replicate sequencing was available, the mean B-cell indel rate across replicates was used. Plots were generated with ggplot2, ggpubr, ggrepel, and scales, and figures were finalized in Adobe Illustrator 2025, with selected elements exported from Biorender.org. Data summaries are presented as boxplots, violin plots, or mean ± SD, as appropriate; specific conventions are noted in figure legends.

### Study approval and Ethics Statement

The patient and her parents provided written informed consent for the collection, analysis, and publication of their medical and genetic data. The study was conducted in accordance with the Declaration of Helsinki and current Danish legislation. The study was also registered in the internal record of research projects at Region Midtjylland (Case No. 1-16-02-238-23).

B cells were purified from buffy coats, obtained through the blood bank at Aarhus University Hospital, Denmark, from anonymized healthy volunteers, in accordance with the Danish Health Care Act (Sundhedsloven) at the time.

## Abbreviations

Aa: Amino-acid residue
Ab: Antibody
AD: Autosomal-dominant
AICDA/AID: Activation-induced cytidine deaminase
AR: Autosomal-recessive
BCR: B-cell receptor
BER: Base-excision repair
CSR: Class-switch recombination
CTRL: Control
DSB: Double-strand break
GC: Germinal center
HIGM2: Hyper-IgM syndrome type 2
hnRNP: heterogenous ribonucleoprotein
Ig: Immunoglobulin
iGB: Inducible germinal center B cell
MBC: Memory B cell
MMR: Mismatch repair
nB: Naïve B cell
NES: Nuclear export signal
PC: Plasma cell
PT: Patient
SHM: Somatic hypermutation
TRIFMA: time-resolved immunofluorometric assay
WT: Wild Type
WGS: Whole-genome sequencing.

## Author contributions

Conceptualization, KG, ABR, ETD, LW and SED; Methodology, ABR, KG, ETD, JMBJ, LJ, JGM and SED; Formal analysis, KG, ABR; Investigation, KG, ABR, ETD, JMBJ, LJ, KM, JHW, LL, and SED; Clinical patient care, CSL; Writing – Original Draft, KG, ABR and SED; Writing – Review & Editing, KG, ABR, JMBJ and SED; Visualization, KG and ABR; Supervision, SED; Project Administration, KG, ABR, and SED; Funding Acquisition, ABR, JMBJ, JGM and SED.

## Acknowledgements

This work was funded by grants from the Independent Research Fund Denmark to JGM (9039-00173B) and SED (Sapere Aude Research Leader grant; 9060-00038). Additional funding was provided by Aase og Ejnar Danielsens Fond and the NEYE foundation. We are grateful to the FACS Core at AU for expert assistance with flow cytometry experiments. Furthermore, we thank Emil Aagaard Thompson for nucleofection of THP-1 cells for testing *AICDA*-targeting gRNAs.

## References

1. Muramatsu M, Kinoshita K, Fagarasan S, Yamada S, Shinkai Y, Honjo T. Class switch recombination and hypermutation require activation-induced cytidine deaminase (AID), a potential RNA editing enzyme. Cell. 2000;102(5):553–63.

2. Vincent-Fabert C, Fiancette R, Pinaud E, Truffinet Vr, Cogné N, Cogné M, et al. Genomic deletion of the whole IgH 3’ regulatory region (hs3a, hs1,2, hs3b, and hs4) dramatically affects class switch recombination and Ig secretion to all isotypes. Blood. 2010;116 11:1895–8.

3. Sabouri S, Kobayashi M, Begum NA, Xu J, Hirota K, Honjo T. C-terminal region of activation-induced cytidine deaminase (AID) is required for efficient class switch recombination and gene conversion. Proceedings of the National Academy of Sciences. 2014;111:2253–8.

4. Hu W, Begum NA, Mondal S, Stanlie A, Honjo T. Identification of DNA cleavage- and recombination-specific hnRNP cofactors for activation-induced cytidine deaminase. Proceedings of the National Academy of Sciences. 2015;112:5791–6.

5. Petersen-Mahrt SK, Harris RS, Neuberger MS. AID mutates E. coli suggesting a DNA deamination mechanism for antibody diversification. Nature. 2002;418(6893):99–103.

6. Chaudhuri J, Tian M, Khuong C, Chua K, Pinaud E, Alt FW. Transcription-targeted DNA deamination by the AID antibody diversification enzyme. Nature. 2003;422(6933):726–30.

7. Bransteitter R, Pham P, Scharff MD, Goodman MF. Activation-induced cytidine deaminase deaminates deoxycytidine on single-stranded DNA but requires the action of RNase. Proc Natl Acad Sci U S A. 2003;100(7):4102–7.

8. Rada C, Di Noia JM, Neuberger MS. Mismatch recognition and uracil excision provide complementary paths to both Ig switching and the A/T-focused phase of somatic mutation. Molecular cell. 2004;16 2:163–71.

9. Delbos F, Aoufouchi S, Faili A, Weill JC, Reynaud CA. DNA polymerase eta is the sole contributor of A/T modifications during immunoglobulin gene hypermutation in the mouse. J Exp Med. 2007;204(1):17–23.

10. Zuo T, Gautam A, Saghaei S, Khobragade SN, Ahmed R, Mahdavinia A, et al. Somatic hypermutation generates antibody specificities beyond the primary repertoire. Immunity. 2025;58(6):1396–410.e7.

11. Revy P, Muto T, Levy Y, Geissmann F, Plebani A, Sanal O, et al. Activation-induced cytidine deaminase (AID) deficiency causes the autosomal recessive form of the Hyper-IgM syndrome (HIGM2). Cell. 2000;102(5):565–75.

12. Yazdani R, Fekrvand S, Shahkarami S, Azizi G, Moazzami B, Abolhassani H, et al. The hyper IgM syndromes: Epidemiology, pathogenesis, clinical manifestations, diagnosis and management. Clinical immunology. 2019;198:19–30.

13. Poli MC, Aksentijevich I, Bousfiha AA, Cunningham-Rundles C, Hambleton S, Klein C, et al. Human inborn errors of immunity: 2024 update on the classification from the International Union of Immunological Societies Expert Committee. Journal of Human Immunity. 2025;1(1).

14. Quartier P, Bustamante J, Sanal O, Plebani A, Debré M, Deville A, et al. Clinical, immunologic and genetic analysis of 29 patients with autosomal recessive hyper-IgM syndrome due to Activation-Induced Cytidine Deaminase deficiency. Clin Immunol. 2004;110(1):22–9.

15. Kasahara Y, Kaneko H, Fukao T, Terada T, Asano T, Kasahara K, et al. Hyper-IgM syndrome with putative dominant negative mutation in activation-induced cytidine deaminase. The Journal of allergy and clinical immunology. 2003;112 4:755–60.

16. Durandy A, Taubenheim N, Péron S, Fischer A. Pathophysiology of B-cell intrinsic immunoglobulin class switch recombination deficiencies. Advances in immunology. 2007;94:275–306.

17. Mina ED, Jackson KJ, Crawford AJI, Faulks ML, Pathmanandavel K, Acquarola N, et al. A Novel Heterozygous Variant in AICDA Impairs Ig Class Switching and Somatic Hypermutation in Human B Cells and is Associated with Autosomal Dominant HIGM2 Syndrome. Journal of Clinical Immunology. 2024;44.

18. Fadlallah J, Chentout L, Boisson B, Pouliet A, Masson C, Morin F, et al. From Dysgammaglobulinemia to Autosomal-Dominant Activation-Induced Cytidine Deaminase Deficiency: Unraveling an Inherited Immunodeficiency after 50 Years. The Journal of pediatrics. 2020.

19. Kacar M, Shrimpton JK, Jassam M, Mistry A, Arumugakani G, Grammatikos A, et al. Hyper-IgM syndrome resulting from heterozygous AICDA variants: A European first? Scand J Immunol. 2022;96(1):e13155.

20. Kuipers C, Hennessey L, Hartog NL. Expanding clinical phenotype of truncating activation-induced cytidine deaminase variants. Ann Allergy Asthma Immunol. 2025;135(4):454–5.

21. Barreto V, Reina-San-Martin B, Ramiro AR, McBride KM, Nussenzweig MC. C-terminal deletion of AID uncouples class switch recombination from somatic hypermutation and gene conversion. Mol Cell. 2003;12(2):501–8.

22. Ta VBT, Nagaoka H, Catalan N, Durandy A, Fischer A, Imai K, et al. AID mutant analyses indicate requirement for class-switch-specific cofactors. Nature Immunology. 2003;4:843–8.

23. Shinkura R, Ito S, Begum NA, Nagaoka H, Muramatsu M, Kinoshita K, et al. Separate domains of AID are required for somatic hypermutation and class-switch recombination. Nature Immunology. 2004;5:707–12.

24. Geisberger R, Rada C, Neuberger MS. The stability of AID and its function in class-switching are critically sensitive to the identity of its nuclear-export sequence. Proc Natl Acad Sci U S A. 2009;106(16):6736–41.

25. Doi T, Kato LM, Ito S, Shinkura R, Wei M, Nagaoka H, et al. The C-terminal region of activation-induced cytidine deaminase is responsible for a recombination function other than DNA cleavage in class switch recombination. Proceedings of the National Academy of Sciences. 2009;106:2758–63.

26. Zahn A, Eranki AK, Patenaude A-M, Methot SP, Fifield H, Cortizas EM, et al. Activation induced deaminase C-terminal domain links DNA breaks to end protection and repair during class switch recombination. Proceedings of the National Academy of Sciences. 2014;111:E988–E97.

27. Ranjit S, Khair L, Linehan EK, Ucher AJ, Chakrabarti M, Schrader CE, et al. AID Binds Cooperatively with UNG and Msh2-Msh6 to Ig Switch Regions Dependent upon the AID C Terminus. The Journal of Immunology. 2011;187:2464–75.

28. Refaat AM, Nakata M, Husain A, Kosako H, Honjo T, Begum NA. HNRNPU facilitates antibody class-switch recombination through C-NHEJ promotion and R-loop suppression. Cell reports. 2023;42 3:112284.

29. Mondal S, Begum NA, Hu W, Honjo T. Functional requirements of AID’s higher order structures and their interaction with RNA-binding proteins. Proceedings of the National Academy of Sciences. 2016;113:E1545–E54.

30. Ito S, Nagaoka H, Shinkura R, Begum N, Muramatsu M, Nakata M, et al. Activation-induced cytidine deaminase shuttles between nucleus and cytoplasm like apolipoprotein B mRNA editing catalytic polypeptide 1. Proceedings of the National Academy of Sciences. 2004;101(7):1975–80.

31. McBride KM, Barreto VM, Ramiro AR, Stavropoulos P, Nussenzweig MC. Somatic Hypermutation Is Limited by CRM1-dependent Nuclear Export of Activation-induced Deaminase. The Journal of Experimental Medicine. 2004;199:1235–44.

32. Patenaude A-M, Orthwein A, Hu Y, Campo VA, Kavli B, Buschiazzo A, et al. Active nuclear import and cytoplasmic retention of activation-induced deaminase. Nature Structural &Molecular Biology. 2009;16:517–27.

33. Methot SP, Litzler LC, Trajtenberg F, Zahn A, Robert F, Pelletier J, et al. Consecutive interactions with HSP90 and eEF1A underlie a functional maturation and storage pathway of AID in the cytoplasm. The Journal of Experimental Medicine. 2015;212:581–96.

34. Pasqualucci L, Guglielmino R, Houldsworth J, Mohr J, Aoufouchi S, Polakiewicz RD, et al. Expression of the AID protein in normal and neoplastic B cells. Blood. 2004;104 10:3318–25.

35. Al Ismail A, Husain A, Kobayashi M, Honjo T, Begum NA. Depletion of recombination-specific cofactors by the C-terminal mutant of the activation-induced cytidine deaminase causes the dominant negative effect on class switch recombination. International Immunology. 2017;29:525–37.

36. Xie X, Gan T, Rao B, Zhang W, Panchakshari RA, Yang D, et al. C-terminal deletion-induced condensation sequesters AID from IgH targets in immunodeficiency. EMBO J. 2022;41(11):e109324.

37. Ucher AJ, Ranjit S, Kadungure T, Linehan EK, Khair L, Xie E, et al. Mismatch Repair Proteins and AID Activity Are Required for the Dominant Negative Function of C-Terminally Deleted AID in Class Switching. The Journal of Immunology. 2014;193:1440–50.

38. Imai K, Zhu Y, Revy P, Morio T, Mizutani S, Fischer A, et al. Analysis of class switch recombination and somatic hypermutation in patients affected with autosomal dominant hyper-IgM syndrome type 2. Clinical immunology. 2005;115 3:277–85.

39. Cantaert T, Schickel J-N, Bannock JM, Ng Y-s, Massad C, Delmotte FR, et al. Decreased somatic hypermutation induces an impaired peripheral B cell tolerance checkpoint. The Journal of clinical investigation. 2016;126 11:4289–302.

40. Wang P, Wang K. PubMind: literature-based genetic variant extraction and functional annotation using large language models. Nat Commun. 2026;17(1).

41. Dickerson SK, Market E, Besmer E, Papavasiliou FN. AID Mediates Hypermutation by Deaminating Single Stranded DNA. The Journal of Experimental Medicine. 2003;197: 1291 - 6.

42. Pham P, Afif SA, Shimoda M, Maeda K, Sakaguchi N, Pedersen LC, et al. Structural analysis of the activation-induced deoxycytidine deaminase required in immunoglobulin diversification. DNA repair. 2016;43:48–56.

43. Xu Z, Zan H, Pone EJ, Mai T, Casali P. Immunoglobulin class-switch DNA recombination: induction, targeting and beyond. Nat Rev Immunol. 2012;12(7):517–31.

44. Larijani M, Petrov AP, Kolenchenko O, Berru M, Krylov SN, Martin A. AID Associates with Single-Stranded DNA with High Affinity and a Long Complex Half-Life in a Sequence-Independent Manner. Molecular and Cellular Biology. 2006;27:20–30.

45. Rogozin IB, Diaz M. Cutting Edge: DGYW/WRCH Is a Better Predictor of Mutability at G:C Bases in Ig Hypermutation Than the Widely Accepted RGYW/WRCY Motif and Probably Reflects a Two-Step Activation-Induced Cytidine Deaminase-Triggered Process. The Journal of Immunology. 2004;172:3382–4.

46. Qiao Q, Wang L, Meng F-L, Hwang JK, Alt FW, Wu H. AID Recognizes Structured DNA for Class Switch Recombination. Molecular cell. 2017;67 3:361–73.e4.

47. Yamane A, Resch W, Kuo N, Kuchen S, Li Z, Sun HW, et al. Deep-sequencing identification of the genomic targets of the cytidine deaminase AID and its cofactor RPA in B lymphocytes. Nat Immunol. 2011;12(1):62–9.

48. Delker RK, Zhou Y, Strikoudis A, Stebbins CE, Papavasiliou FN. Solubility-based genetic screen identifies RING finger protein 126 as an E3 ligase for activation-induced cytidine deaminase. Proc Natl Acad Sci U S A. 2013;110(3):1029–34.

49. Xu Z, Fulop Z, Wu G, Pone EJ, Zhang J, Mai T, et al. 14-3-3 adaptor proteins recruit AID to 5’-AGCT-3’-rich switch regions for class switch recombination. Nat Struct Mol Biol. 2010;17(9):1124–35.

50. Cheng J, Novati G, Pan J, Bycroft C, Žemgulytė A, Applebaum T, et al. Accurate proteome-wide missense variant effect prediction with AlphaMissense. Science. 2023;381(6664):eadg7492.

51. Hao Y, Stuart T, Kowalski MH, Choudhary SK, Hoffman PJ, Hartman A, et al. Dictionary learning for integrative, multimodal and scalable single-cell analysis. Nature Biotechnology. 2022;42:293–304.

52. Gupta NT, Heiden JAV, Uduman M, Gadala-Maria D, Yaari G, Kleinstein SH. Change-O: a toolkit for analyzing large-scale B cell immunoglobulin repertoire sequencing data. Bioinformatics. 2015;31 20:3356–8.

53. Rovsing AB, Green K, Jensen L, Nielsen IH, Mikkelsen JG, Degn SE. Multiparametric Optimization of Human Primary B-Cell Cultures Using Design of Experiments. Scand J Immunol. 2025;102(2):e70043.

54. Muramatsu M, Sankaranand VS, Anant S, Sugai M, Kinoshita K, Davidson NO, et al. Specific expression of activation-induced cytidine deaminase (AID), a novel member of the RNA-editing deaminase family in germinal center B cells. J Biol Chem. 1999;274(26):18470–6.

55. Cattoretti G, Büttner M, Shaknovich R, Kremmer E, Alobeid B, Niedobitek G. Nuclear and cytoplasmic AID in extrafollicular and germinal center B cells. Blood. 2006;107(10):3967–75.

56. Wu CM, Roth TL, Baglaenko Y, Ferri DM, Brauer P, Zuniga-Pflucker JC, et al. Genetic engineering in primary human B cells with CRISPR-Cas9 ribonucleoproteins. J Immunol Methods. 2018;457:33–40.

57. Aoufouchi S, Faili A, Zober C, D’Orlando O, Weller S, Weill JC, et al. Proteasomal degradation restricts the nuclear lifespan of AID. J Exp Med. 2008;205(6):1357–68.

58. Uchimura Y, Barton LF, Rada C, Neuberger MS. REG-γ associates with and modulates the abundance of nuclear activation-induced deaminase. J Exp Med. 2011;208(12):2385–91.

59. Al-Saud B, Al-Jomaie M, Al-Ghonaium A, Al-Ahmari A, Al-Mousa H, Al-Muhsen S, et al. Haematopoietic stem cell transplant for hyper-IgM syndrome due to CD40 defects: a single-centre experience. Bone Marrow Transplant. 2019;54(1):63–7.

60. Ferrua F, Galimberti S, Courteille V, Slatter MA, Booth C, Moshous D, et al. Hematopoietic stem cell transplantation for CD40 ligand deficiency: Results from an EBMT/ESID-IEWP-SCETIDE-PIDTC study. J Allergy Clin Immunol. 2019;143(6):2238–53.

61. Tangye SG, Al-Herz W, Bousfiha A, Chatila T, Cunningham-Rundles C, Etzioni A, et al. Correction to: Human Inborn Errors of Immunity: 2019 Update on the Classification from the International Union of Immunological Societies Expert Committee. J Clin Immunol. 2020;40(1):65.

62. Jumper J, Evans R, Pritzel A, Green T, Figurnov M, Ronneberger O, et al. Highly accurate protein structure prediction with AlphaFold. Nature. 2021;596(7873):583–9.

63. Tordai H, Torres O, Csepi M, Padányi R, Lukács GL, Hegedűs T. Analysis of AlphaMissense data in different protein groups and structural context. Sci Data. 2024;11(1):495.

64. Laine E, Karami Y, Carbone A. GEMME: A Simple and Fast Global Epistatic Model Predicting Mutational Effects. Mol Biol Evol. 2019;36(11):2604–19.

65. Abakarova M, Freiberger MI, Liehrmann A, Rera M, Laine E. Proteome-wide Prediction of the Functional Impact of Missense Variants with ProteoCast. bioRxiv. 2025:2025.02.09.637326.

66. Sievers F, Wilm A, Dineen D, Gibson TJ, Karplus K, Li W, et al. Fast, scalable generation of high-quality protein multiple sequence alignments using Clustal Omega. Mol Syst Biol. 2011;7:539.

67. Rentzsch P, Witten D, Cooper GM, Shendure J, Kircher M. CADD: predicting the deleteriousness of variants throughout the human genome. Nucleic Acids Res. 2019;47(D1):D886–d94.

68. McGinnis CS, Murrow LM, Gartner ZJ. DoubletFinder: Doublet detection in single-cell RNA sequencing data using artificial nearest neighbors.bioRxiv. 2018.

69. Zappia L, Oshlack A. Clustering trees: a visualization for evaluating clusterings at multiple resolutions. GigaScience. 2018;7.

70. Magoč T, Salzberg SL. FLASH: fast length adjustment of short reads to improve genome assemblies. Bioinformatics. 2011;27(21):2957–63.

71. Clement K, Rees H, Canver MC, Gehrke JM, Farouni R, Hsu JY, et al. CRISPResso2 provides accurate and rapid genome editing sequence analysis. Nat Biotechnol. 2019;37(3):224–6.

72. Xu S, Hu E, Cai Y, Xie Z, Luo X, Zhan L, et al. Using clusterProfiler to characterize multiomics data. Nat Protoc. 2024;19(11):3292–320.

73. Wu T, Hu E, Xu S, Chen M, Guo P, Dai Z, et al. clusterProfiler 4.0: A universal enrichment tool for interpreting omics data. The Innovation. 2021;2.

